# eATP/P2X7R axis drives nanoparticle induced neutrophil recruitment in the pulmonary microcirculation

**DOI:** 10.1101/2024.03.11.584398

**Authors:** Chenxi Li, Qiongliang Liu, Lianyong Han, Roland Immler, Birgit Rathkolb, Judith Secklehner, Martin Hrabe de Angelis, Ali Önder Yildirim, Annette Nicke, Leo M. Carlin, Markus Sperandio, Tobias Stoeger, Markus Rehberg

**Affiliations:** Institute of Lung Health and Immunity (LHI), Comprehensive Pneumology Center (CPC), Helmholtz Center Munich; Member of the German Center for Lung Research (DZL), Munich, Germany; Department of Thoracic Surgery, Shanghai General Hospital, Shanghai Jiao Tong University School of Medicine, Shanghai, 200080, China; Walter Brendel Centre of Experimental Medicine, Biomedical Center, Institute of Cardiovascular Physiology and Pathophysiology, Ludwig-Maximilians-Universität München, Planegg-Martinsried, Germany; Institute of Experimental Genetics and German Mouse Clinic, Helmholtz Zentrum München, Neuherberg, Germany; Institute of Experimental Animal breeding and Biotechnology, Ludwig-Maximilians-Universität München, Munich, Germany; Chair of Experimental Genetics, TUM School of Life Sciences, Technische Universität München, Freising, Germany; Cancer Research UK Scotland Institute, Glasgow, UK; School of Cancer Sciences, University of Glasgow, Glasgow, UK; Walther Straub Institute of Pharmacology and Toxicology, Faculty of Medicine, Ludwig-Maximilians-Universität München, Munich, Germany

## Abstract

Exposure to nanoparticles (NPs) is frequently associated with adverse cardiovascular effects. In contrast, NPs in nanomedicine hold great promise for precise lung-specific drug delivery, especially considering the extensive pulmonary capillary network that facilitates interactions with bloodstream-suspended particles. Therefore, exact knowledge about interactions and effects of engineered NPs with the pulmonary microcirculation are instrumental for future application of this technology in patients. To unravel the real-time dynamics of intravenously delivered NPs and their effects in the pulmonary microvasculature, we employed intravital microscopy of the mouse lung. PEG amine-modified quantum dots (aQDs) with a low potential for biomolecule and cell interactions and carboxyl-modified quantum dots (cQDs) with a high interaction potential were used, representing two different NP subtypes.

Only aQDs triggered rapid neutrophil recruitment in microvessels and their subsequent recruitment to the alveolar space. Application of specific inhibitors revealed that the aQDs induced neutrophil recruitment was linked to cellular degranulation, TNF-α, and DAMP release into the circulation, particularly extracellular ATP (eATP). Stimulation of the ATP-gated P2X7R induced the expression of E-selectin on microvascular endothelium with the subsequent E-selectin depended neutrophilic immune response. Leukocyte integrins (LFA-1 and MAC-1) mediated adhesion and reduction in neutrophil crawling velocity on the vascular surface.

In summary, this study unravels the complex cascade of neutrophil recruitment during NP-induced sterile inflammation. Thereby we demonstrate novel adverse effects for NPs in the pulmonary microcirculation and provide critical insights for optimizing NP-based drug delivery and therapeutic intervention strategies, to ensure their efficacy and safety in clinical applications.

**Graphical Abstract:** 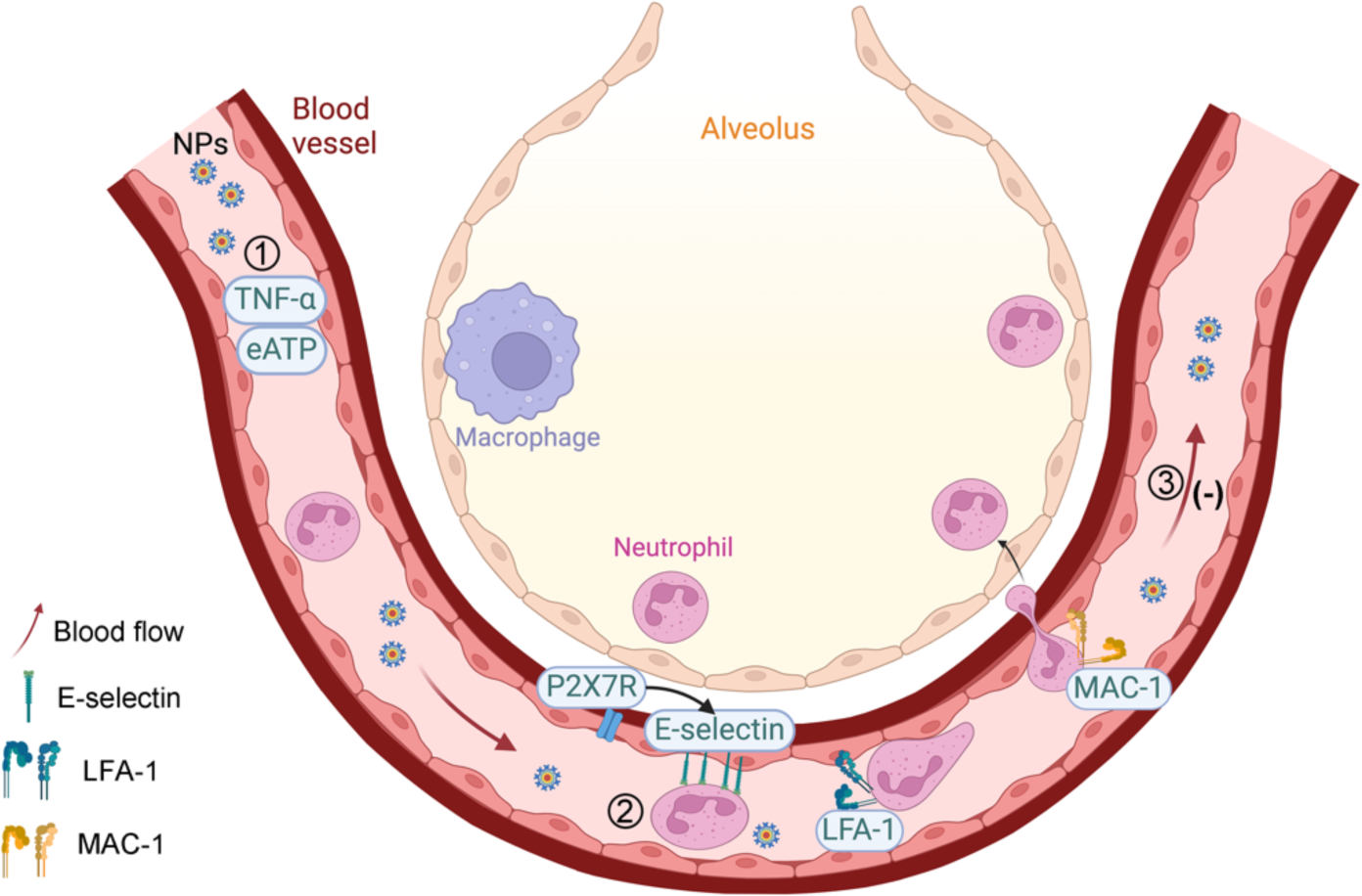

## Introduction

Nanoparticles (NPs) have been widely applied in nanomedicine for drug delivery and as imaging agents. For administration, intravenous injection is a common approach in medical diagnosis and disease treatment (*1*). At the same time, the increasing applications of nanotechnology are exposing individuals to various nanomaterials, raising concerns about potential risk for lung injury, cardiovascular diseases, neurological diseases, auto-immune diseases, and cancer (*2–4*). After inhalation, NPs below 20 nm can evade surveillance by alveolar macrophages that enables them to breach the air-blood barrier, facilitating their translocation into the bloodstream and secondary organs (*2, 5–8*). The lung, characterized by its abundant pulmonary capillary bed, serves as an important surface for interactions with NPs in the bloodstream (*9–11*). The defense niche within the pulmonary microvasculature is distinctive from most extrapulmonary tissues. Neutrophils in the lung, as opposed to resident vascular macrophages, patrol the pulmonary microcirculation as marginated pool, protecting from invading pathogens (*12–14*). The low blood flow velocity in pulmonary microvessels in combination with their small diameter (*15, 16*) could contribute to a prolonged retention of NPs in the pulmonary microcirculations by increased probability of interactions with microvessels and neutrophils.

Quantum dots (QDs), which are fluorescent inorganic semiconductor crystals, represent some of the earliest and most commonly used nanomaterials (*17*). Their exceptional fluorescent properties make them well-suited for a variety of imaging techniques and drug delivery applications, contributing to theragnostic approaches (*17, 18*). Surface modifications of NPs have demonstrated the ability to modulate inflammation and cancer (*19–21*) and these modifications also influence organ distribution and immune responses (*22–24*). However, our understanding of NPs in the bloodstream and their interactions with and effects on immune cells in pulmonary microvessels remains limited (*25*). Although intravital microscopy (IVM) has been used extensively in the study of NP effects and interactions in the microcirculation e.g. in skeletal and hepatic tissues (*26–29*), its application to the pulmonary microcirculation has only recently been addressed. Lung intravital microcopy (L-IVM) enables real-time analysis of dynamic processes and cellular-level imaging in the alveolar region of the murine lung (*30*). This advanced technique has been employed to visualize the pulmonary endothelial surface layer and alveolar epithelial cells (*31, 32*). In the present work, we utilize QD nanoparticles with two different surface modifications: amine-PEG-QDs (aQDs) and carboxylated QDs (cQDs). In our previous work conducted in skeletal muscle, both QD-types did not induce leukocyte recruitment in microvessels of healthy mice, however leukocyte recruitment was induced after trauma induced macrophage activation, solely by cQDs, whereas only application of aQDs aggravated neutrophil responses after ischemia-reperfusion injury in skeletal muscle tissue (*24, 33*).

Upon intravascular delivery of NPs into the bloodstream, interactions with the endothelium and other immune cells might initiate neutrophil recruitment. Endothelial activation is categorized into two types. Type I activation, mediated by ligands like histamine H1 receptors, leads to a transient increase in cytosolic Ca^2+^ levels. This activation is independent of new gene expression and protein synthesis, spontaneously declining within 10–20 minutes. Type II activation is mediated by pro-inflammatory cytokines such as TNF-α and IL-1, lasting for hours to days (*34*). By cQDs induced leukocyte recruitment in skeletal muscle postcapillary venules is effectively inhibited by prior application of cromolyn, an inhibitor of cellular degranulation (*33*). Cromolyn is known to primarily act on mast cells but also on (alveolar) macrophages (*35*), both of which are recognized for releasing histamine and a range of mediators that induce the inflammatory process. Activated endothelial cells, alongside priming agents like pathogen or damage associated molecular patterns (PAMPs, DAMPs), including ATP, and cytokines, stimulate neutrophils, initiating them from a latent state. These priming agents swiftly induce integrin/selectin expression and cellular degranulation, with ATP being the most rapid initiator of this priming process (*36*). Inflammation significantly impacts blood flow in the pulmonary microvessels (*37*). The reduced blood flow creates an opportunity for neutrophils to engage with endothelial cells, potentially leading to their priming and increased responsiveness (*38*). Thus blood borne neutrophils are mobilized to the site of inflammation within minutes, without cytokine synthesis, for functions like phagocytosis, degranulation, or NET release to eliminate pathogens (*12, 39*).

While nanomedical particles targeting the pulmonary microcirculation hold significant promise for enhanced drug delivery and treatment efficacy (*40*), our understanding of the intricate interactions between NPs and the pulmonary microcirculation is still incomplete, potentially impeding their optimal effectiveness and safety. Consequently, a meticulous assessment of the potential risks and benefits associated with the delivery of nanomedicine to the pulmonary microcirculation is imperative. To this end, we employ QDs as model NPs to comprehensively analyze their dynamics and proinflammatory effects in the pulmonary microcirculation of healthy mice, by means of L-IVM.

## Results

### Interactions of aQDs and cQDs within the pulmonary microcirculation

To test QD interactions and cellular uptake by endothelial cells, human umbilical vein endothelial cells (HUVEC) were incubated with fluorescent cQDs and aQDs (emission 655nm) for 1 hour (Fig. 1 A). cQDs, characterized by strong biomolecule binding as previously shown (*23, 24, 33*), were taken up by HUVEC cells, while aQDs, which exhibit weak biomolecule interactions, showed minimal cellular uptake, in accordance with prior research findings (*23, 24, 33, 41*).

**Figure 1:**
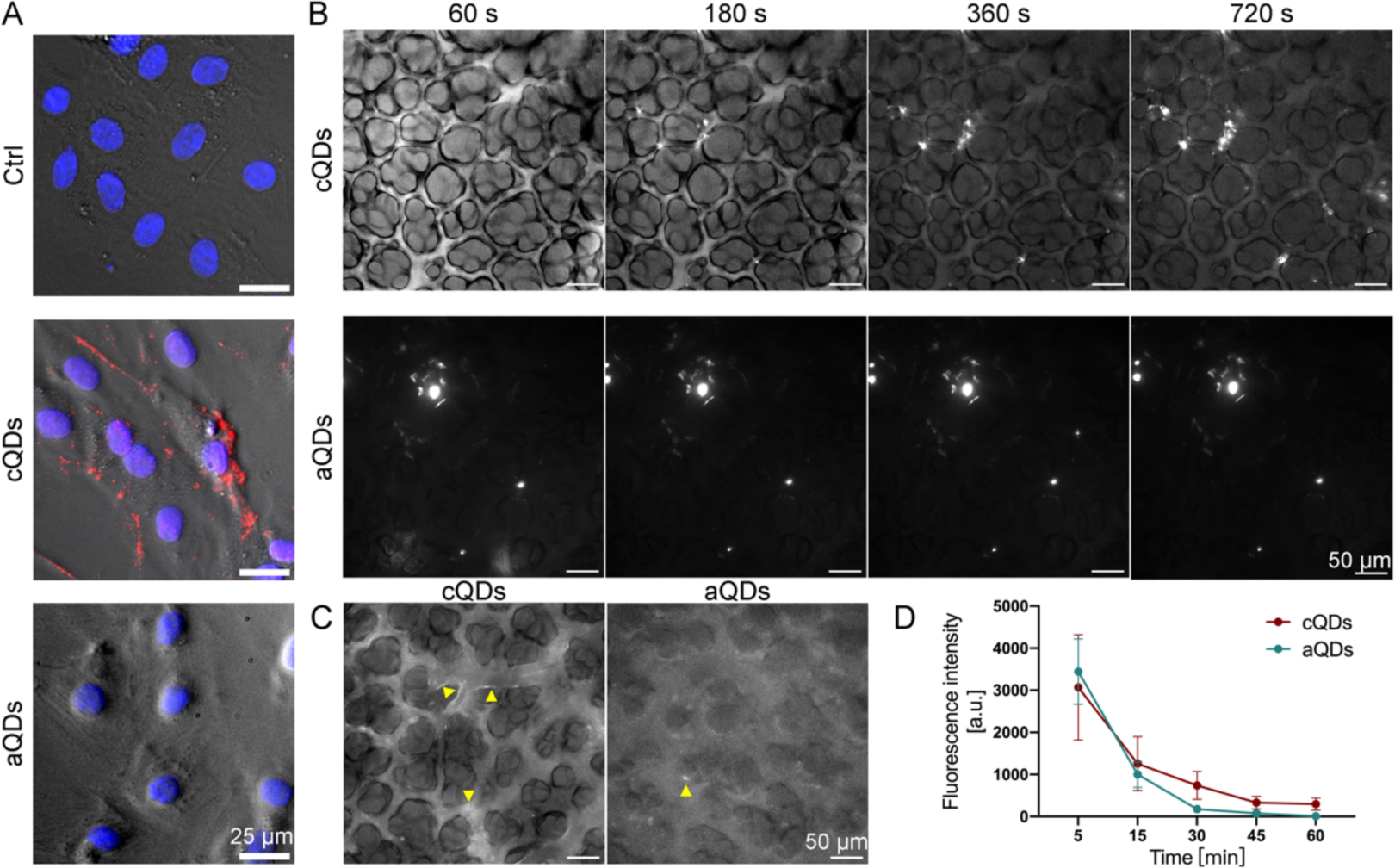
QDs dynamics and localization in vitro and in vivo. (A) The interaction between QDs and endothelial cells. HUVECs were incubated with cQDs and aQDs for 1 h in vitro, followed by fixation with 4 % PFA for IF staining. Phase-contrast images of HUVECs, displayed in grey, show accumulation of cQDs (red), with DAPI-stained nuclei appearing in blue. (Scale bar: 25 μm). (B) Time-lapse imaging reveals the interaction of cQDs with blood vessel walls and the clustering of cQDs and aQDs within pulmonary microvessels via L-IVM. Fluorescence signal (white) solely results from QDs fluorescence, outlining blood vessels and alveolar space. Image brightness in the middle panel was adjusted to display the very bright aQD clusters. (C) Representative images illustrate the pulmonary endothelial uptake of cQDs after 60 minutes whereas only few aQDs spots are present at microvessel walls (both indicated by yellow arrowheads), (Scale bar: 50 µm). (D) Measurement of QD fluorescence intensities in microvessels. Data is shown as mean ± SEM, n = 4 mice/group. Two-way ANOVA test.

To investigate QDs interactions and dynamics in the pulmonary microcirculation by means of L-IVM, aQDs and cQDs (1 pmol/g, respectively) were injected intravenously into mice (Fig. 1 B, C, and D). Both types of QDs could be detected as homogenous fluorescent signal immediately after injection, distributing in the lumen of pulmonary microvessels. cQDs interacted with the blood vessel walls and appeared from the beginning as a lining along the endothelial layer. This localization remained unchanged for 60 min, indicating endothelial cell uptake (Fig. 1 C). aQDs did not line along microvessel walls, but immobile aQDs fluorescent clusters, appeared soon after application and were also stable for 60 min. The fluorescence signals of cQDs as well as aQDs in region of interests covering microvessel segments gradually declined almost to background levels by 60 min (Fig. 1 D), indicating NP clearance from the blood stream.

The lung tissue distribution of QDs one hour after i.v. injection was examined using 3D light sheet fluorescence microscopy (Sup. 1 images & video). Imaging of control lungs displayed clear lung and bronchial contours, outlined by tissue autofluorescence. Mice treated with cQDs after 1 hour exhibited sparse peripheral QD-fluorescence. Conversely, and similar as described by IVM (Fig. 1 B) aQDs were detected throughout the lung, forming aggregates in blood vessels. In conclusion, cQDs and aQDs exhibited different distribution patterns in mouse lungs. Both in L-IVM and 3D light sheet microscopy, we observed that cQDs in pulmonary capillaries were more dispersed and were likely taken up by endothelial cells, wheras aQDs formed some aggregates and remained in the circulation.

### aQDs induce rapid neutrophil recruitment involving alterations in intravascular neutrophil crawling, and a reduction of blood flow velocity

Neutrophil recruitment after cQDs and aQDs application, was assessed through L-IVM (Fig. 2 A). Prior to L-IVM, mice were injected intravenously with Alexa488-labeled anti-Ly6G antibodies which directly labels vascular neutrophils. Images were recorded immediately after surgical insertion of the thoracic window, every 5 to 10 min for 1 hour. As presented in Fig. 2 A and B, i.v. injection of aQDs (1 pmol/g) significantly increased neutrophil numbers already at 20 min upon application, reaching 6.01 ± 0.37/10^4^μm^2^ at 60 min, as compared to neutrophil counts detected in the control group (2.37 ± 0.16/10^4^μm^2^). In contrast, cQDs (1 pmol/g) induced only a very small increase over the observation time (3.46 ± 0.42/10^4^μm^2^ at 60 min). Immunofluorescence staining of neutrophils in lung slices confirmed aQDs’ potent induction of neutrophil recruitment within 1 hour (Sup. 2). Whereas white blood cell (WBC) numbers after both QDs applications remained largely unaffected (Sup. 3 A), aQDs increased neutrophil cell counts and their percentages within WBC and decreased the percentages of monocytes present in the WBC. Interestingly, the application of cQDs for 24 hours induced a decrease in WBC and lymphocyte counts.

**Figure 2.**
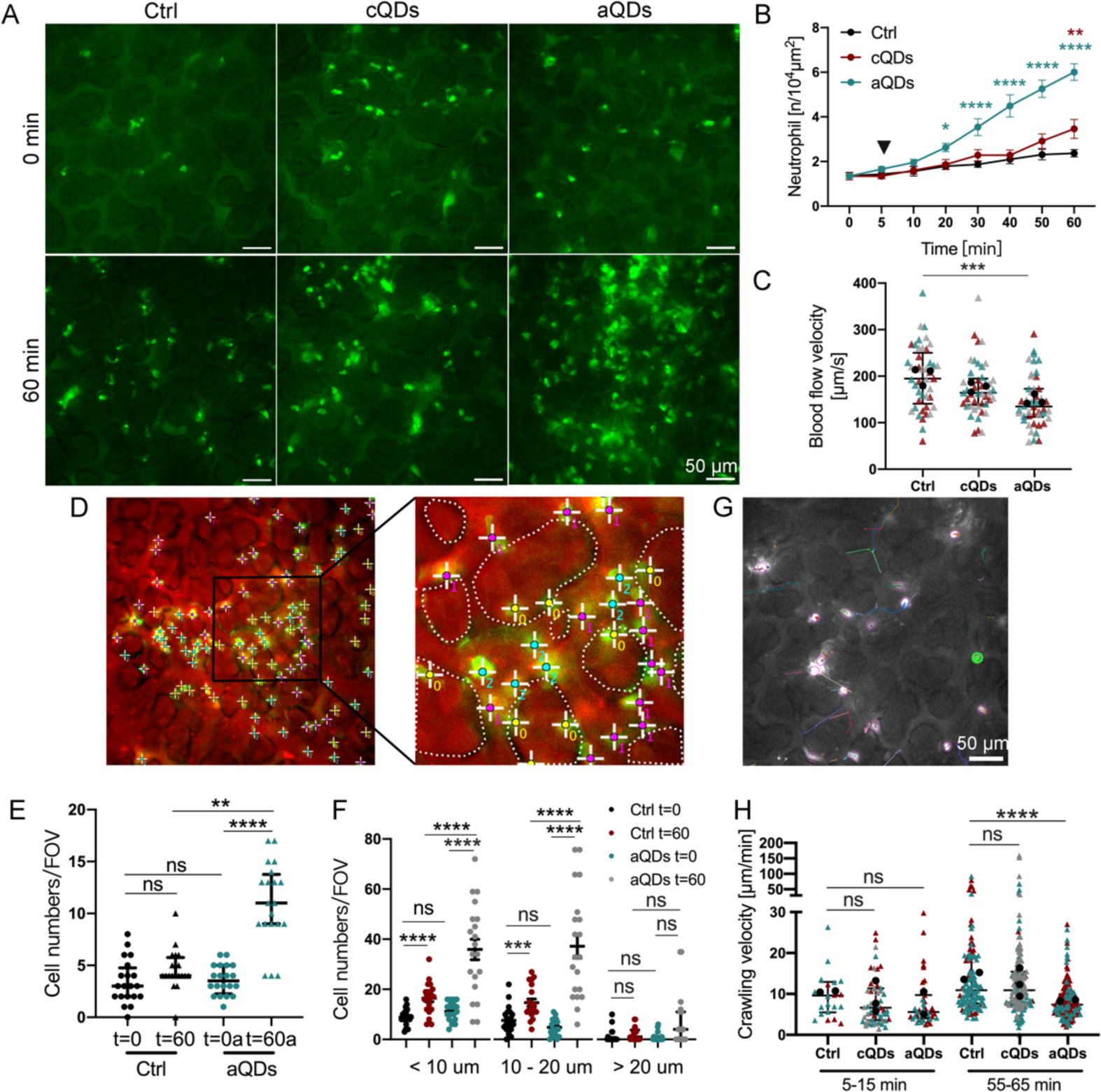
The effect of QDs on neutrophil responses within the pulmonary microcirculation. (A) Representative images depict neutrophils in pulmonary microvessels at 0 min and 60 min after intravenous administration of vehicle (Ctrl), cQDs, and aQDs (1 pmol/g of BW). Neutrophils were directly labeled in vivo by i.v. application of Alexa488-conjugated anti-Ly6G Abs, depicted in bright green points. Autofluorescence outlines alveolar and vascular structures. (Scale bar, 50 μm.) (B) Changes in neutrophil numbers during a 60-min period in mice undergoing QDs application. Data is shown as mean ± SEM, n = 4 mice/group, Two-way ANOVA test, green * indicate significances between aQDs and Ctrl group, red * indicate cQDs and Ctrl group. (C) Blood flow velocity after 60 min in mice undergoing QDs application. Blood velocity was measured by tracking iv. applied fluorescent tracer beads. Data is shown as median (interquartile range, IQR), each measurement is represented by a colored triangle, n = 45 measurements from 3 mice/group, Kruskal-Wallis test. The mean values of the individual mice are shown as black dots. (D) Localization of neutrophils in the alveolar space or microvessels. “0” indicates neutrophils localized in the alveolar space; “1”: neutrophils in vessels smaller than 10 µm; “2”: neutrophils in vessels ranging from 10-20 µm; and “3”: neutrophils in vessels larger than 20 µm. Dotted lines represent alveolar boundaries. Quantification of neutrophils localized in the pulmonary alveolar space (E) and in various-sized microvessels (F). Data is shown as median (IQR) or mean ± SEM, n =20 field of views (FOV) from 4 mice/group, Kruskal-Wallis test or One-way ANOVA test. (G) Representative neutrophil movements over a period of 10 minutes. Neutrophil movements were recorded at 5-second intervals for 10 min using L-IVM. The image displays neutrophil trajectories recorded between minutes 55 and 65 under control conditions, with each color representing the track of an individual neutrophil. (Scale bar: 50 µm). (H) Neutrophil crawling velocities under various applications and time points. Quantification of neutrophil crawling velocities at 5-15 and 55-65 min after vehicle (Ctrl.), aQDs, and cQDs application. Data is shown as median (IQR), each measurement is represented by a colored triangle, n = 22-174 neutrophils analyzed in 2-3 mice/group, Kruskal-Wallis test. The mean values of the individual mice are shown as black dots. * indicates P ≤ 0.05, ** indicates P ≤ 0.01, *** indicates P ≤ 0.001, and **** indicates P ≤ 0.0001.

Neutrophilic inflammation in the lung microvasculature has been shown by L-IVM to be associated with impaired blood flow velocity (*37*). Consequently, we conducted a detailed investigation of the blood velocity of the pulmonary microcirculation, by tracking of i.v. applied fluorescent tracer beads, following QDs application. Baseline blood flow velocity (197.8 μm/s) was reduced significantly to 137.5 μm/s after 1-hour of aQDs application, while cQDs only slightly lowered blood flow to 167.3 μm/s (Fig. 2 C and Sup. 4), thus indicating, that the aQD-induced inflammatory response was accompanied by a reduction in blood flow in the pulmonary microcirculation.

For the precise analysis in which microvessel subtypes neutrophils arrested after aQD induced sterile inflammation, microvessels in L-IVM images at 0 min and 60 min were categorized according to their diameter (< 10 μm, 10-20 μm, > 20 μm, Fig. 2 D). In addition, neutrophil transmigration to the alveolar space was determined. In control mice, most neutrophils were found in vessels < 20 μm (Fig. 2 E and F). Following exposure to aQDs, neutrophil numbers were notably elevated compared to control values, particularly in microvessels smaller than 20 μm, as well as in the alveolar space. This indicates that aQD induced neutrophil recruitment primarily took place in microvessels <20 μm, particularly around alveoli. Notably, neutrophil transmigration to the alveolar space was evident at this early time point after NP injection, resulting in a sustained influx of neutrophils into the alveolar space, as indicated by an increase in neutrophils count at 24 hours in bronchioalveolar lavage fluid (BALF) (Sup. 3 B). Since neutrophils undergo crawling in the lung microvasculature, previously reported for the clearance of bloodstream pathogens (*42*), we conducted an in-depth study on neutrophil motility after i.v. application of aQDs. This investigation involved long-term recordings at 5-second intervals in high-quality image areas (Fig. 2 G and Sup. 5 video), aiming to understand the dynamics of vascular neutrophil behavior in response to nanoparticle exposure. Crawling neutrophils were defined as cells interacting with microvessels for >30 s, while exhibiting movement, according to current literature (*42*). Neutrophil crawling velocities were analyzed in two 10-min intervals (5-15 and 55-65 min, Fig. 2 H). Under control conditions neutrophils crawled at 9.6 μm/min. After QDs injection, the aQDs group exhibited a velocity drop to 5.6 μm/min, comparable to the cQDs group 6.6 μm/min). The aQDs group showed a decreased crawling velocity 7.4 μm/min) after one hour in comparison to the velocities in control and cQDs group (10.9 μm/min and 10.8 μm/min, respectively). The findings indicate that aQDs induced a more persistent inflammatory response, which slowed the motility of crawling neutrophils in pulmonary microvessels, potentially through the induction of specific adhesion molecules.

### Cellular degranulation and TNF-α release is required for the aQD induced immune response

Cromolyn, a cellular degranulation inhibitor, attenuated cQDs-induced leukocyte recruitment in skeletal muscle, implicating mast cell or macrophage participation (*33, 43*). We examined whether cromolyn curb the inflammatory response elicited by aQDs in the pulmonary immune system. Administering 0.2 mg/kg (BW) of cromolyn to mice 30 min prior to aQDs application, significantly reduced aQDs-triggered neutrophil recruitment, (3.04 ± 0.62/10^4^μm^2^ in cromolyn + aQDs group vs. 6.01 ± 0.37/10^4^μm^2^ in aQDs group) (Fig.3 A). Notably, no mast cells, characterized by purple granules stained with toluidine blue, were observed in the peripheral or alveolar regions of the lungs, neither in tissue sections of control mice, nor after 1 hour of aQD application. However, after 24 hours of exposure with aQDs as well as 4 hours after instillation of LPS (which served as a positive inflammatory control), few toluidine blue positive mast cells could be detected near bronchioles (Sup. 6). Alveolar localized mast cells were undetectable at both time points in all conditions. The presence of mast cells in the lung parenchyma has recently been linked to the hygiene status of the mice. Mice housed under ‘specific pathogen-free’ conditions, are in contrast to wild mice, almost devoid of these cells (*44*). Therefore, in our model, immediate involvement of alveolar-localized mast cells in local neutrophil recruitment seems unlikely, yet mast cells in other tissues could contribute, potentially also increasing in number or granularity following QDs exposure. In general, mast cells and macrophages effectively initiate neutrophil recruitment through TNF-α release, early during the inflammatory response (*43, 45, 46*). To test whether TNF-α is involved in aQDs-triggered neutrophil recruitment, we i.v. applied neutralizing anti-TNF-α monoclonal antibodies (mAbs) 30 min before aQDs administration. Anti-TNF-α mAbs significantly reduced basal neutrophil cell counts (t=0 min) during L-IVM, dropping to 0.85 ± 0.13/10^4^μm^2^ in comparison to control values of 1.36 ± 0.16/10^4^μm^2^ (Fig.3 B). Post-aQDs application, neutrophil levels remained below control, reaching 1.83 ± 0.13/10^4^μm^2^ after 60 min. In addition, one hour post L-IVM, cromolyn as well as anti-TNF-α mAbs in aQDs-exposed mice notably increased blood flow velocity to 201.8 μm/s and 231.9 μm/s, respectively (Fig.3 C). The number of neutrophils detected in histology slides confirmed the effects of cromolyn and TNF-α on neutrophil recruitment elicited by aQDs as observed by L-IVM (Fig.3 D). Interestingly, also pretreatment with IgG1 isotype control antibodies diminished to some extend aQDs-induced neutrophil recruitment, albeit less effectively than anti-TNF-α mAbs (Fig.3 B). Previous studies reported that application of IgG can decrease levels of pro-inflammatory cytokines such as IL-1β and IL-6, which are associated with neutrophil recruitment (*47*). Anti-inflammatory properties of IgG1, possibly through C5a pathway blockade (*48*), could explain this observation. Neither application of IgG2a (see Fig. 5 A) nor IgG2b (see Fig. 4 and 5 B) had this effect.

**Figure 3.**
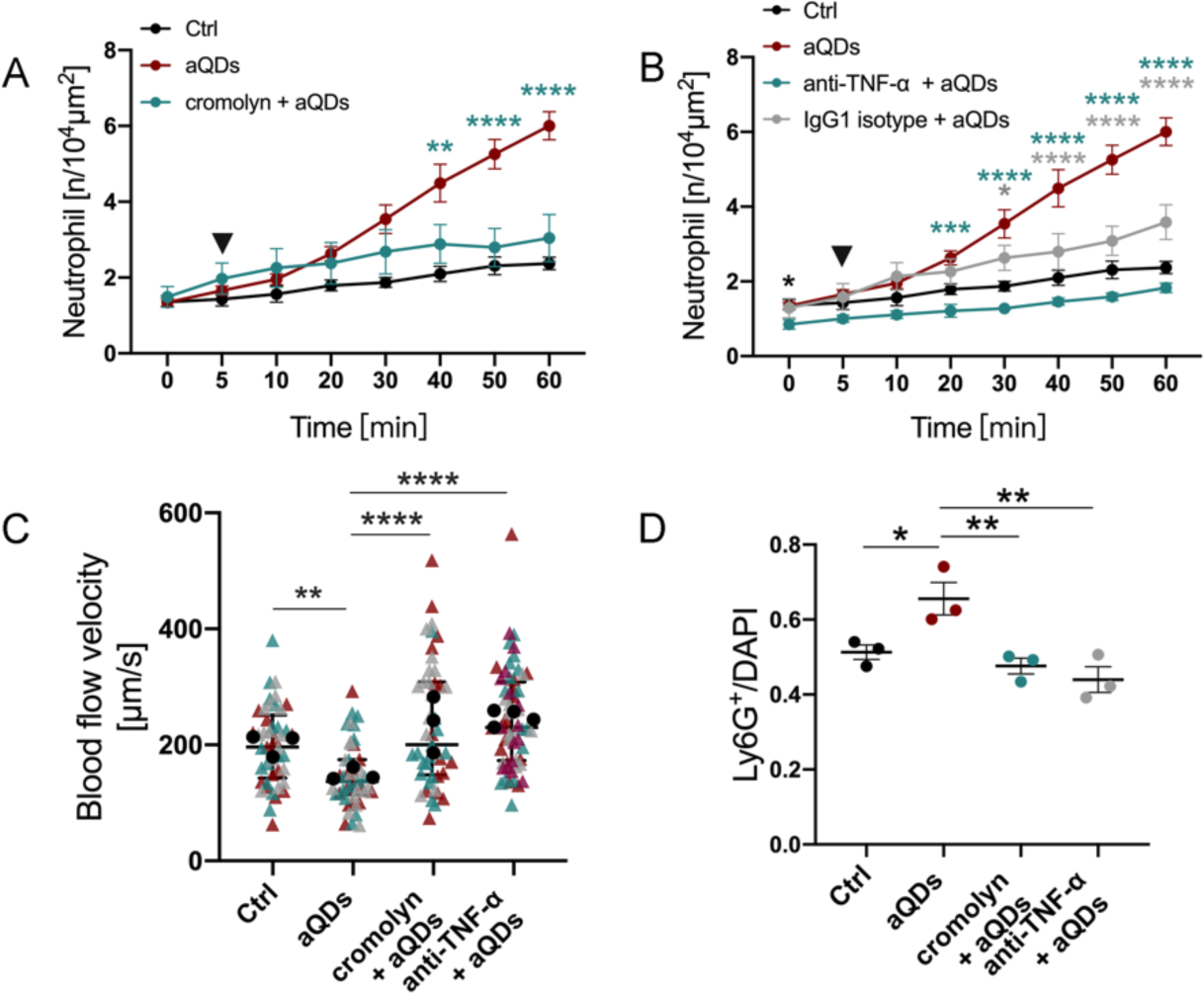
Suppression of aQDs-induced neutrophil recruitment by Cromolyn or TNF-α neutralization in the pulmonary microcirculation. (A) Neutrophil recruitment induced by aQDs was inhibited by Cromolyn. Cromolyn (0.2 mg/kg of BW) was intravenously applied at 30 min prior to L-IVM and aQDs or vehicle were injected at 5 min (arrowhead). Data is shown as mean ± SEM, n = 4 mice/group, Two-way ANOVA test, green stars indicate significances between the cromolyn + aQDs and aQDs groups. (B) anti-TNF-α mAbs diminished neutrophil recruitment induced by aQDs. Anti-TNF-α mAbs (30 μg/mouse) were intravenously applied at 30 min prior to L-IVM. Data is shown as mean ± SEM, n = 4 mice/group, control and aQD neutrophil counts same data as in Fig. 2B, Student’s t-test, black stars indicate significances between anti-TNF-α mAbs + aQDs and Ctrl group; Two-way ANOVA test, green stars indicate significances between anti-TNF-α mAbs + aQDs and aQDs group, grey stars indicate anti-IgG1 mAbs + aQDs and aQDs group. (C) Reduced blood flow velocity after aQDs injection was recovered after pretreatment with cromolyn or anti-TNF-α mAbs. Data is shown as median (IQR), each measurement is represented by a colored triangle, n = 45-60 measurements from 3-4 mice/group, Kruskal-Wallis test. The mean values of the individual mice are shown as black dots. (D) Quantification of Ly6G^+^ neutrophils immune-stained tissue samples. Data is shown as mean ± SEM, n = 3 mice (6 FOV) /group, One-way ANOVA test. * indicates P ≤ 0.05, ** indicates P ≤ 0.01, *** indicates P ≤ 0.001, and **** indicates P ≤ 0.0001.

**Figure 4.**
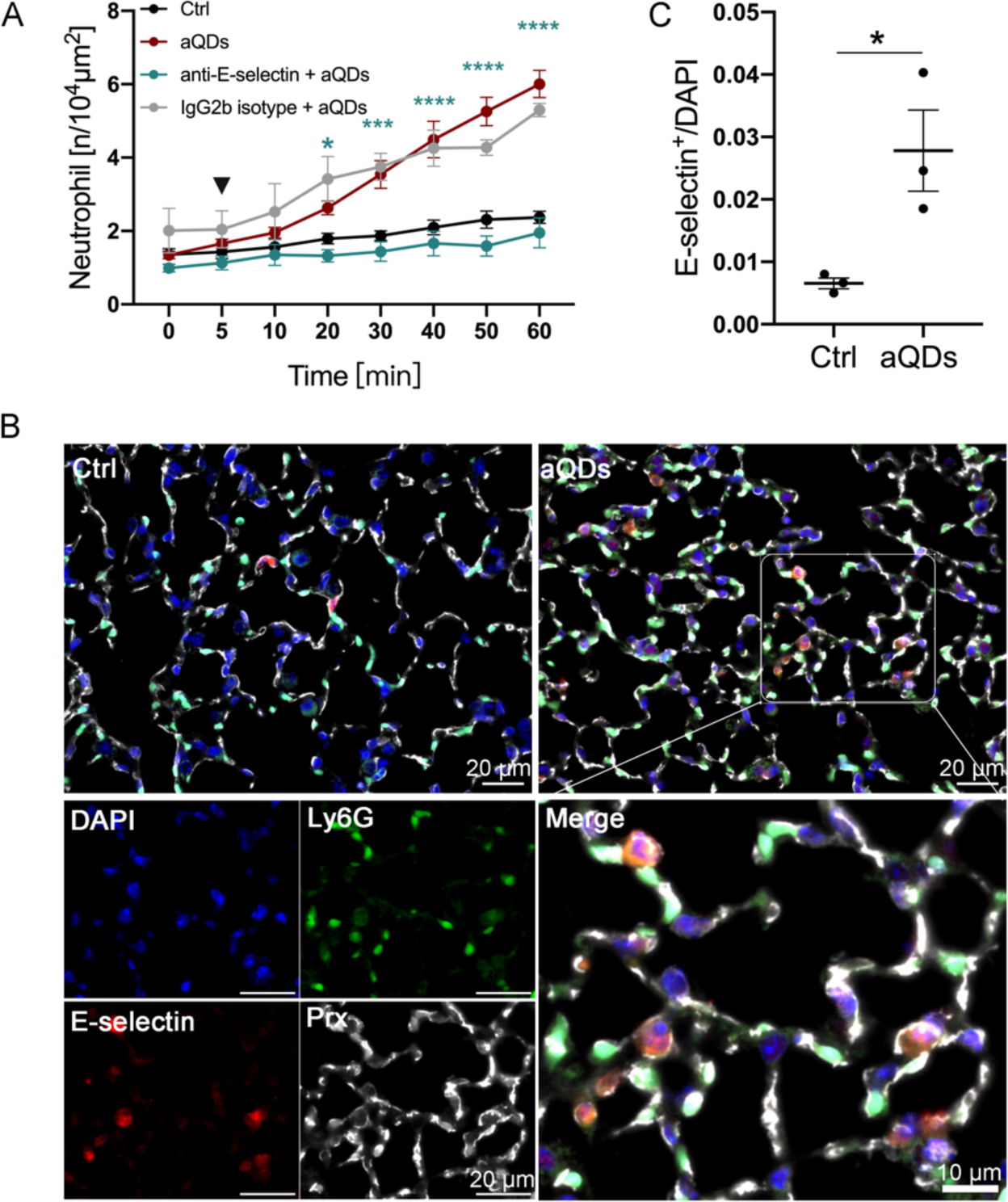
aQDs-induced neutrophil recruitment is mediated by E-selectin. (A) Quantification of recruited neutrophil numbers over time. The mice were pre-treated intravenously with anti-E-selectin mAb 9A9 (20 μg/mouse) for 30 min followed by application of aQDs (arrowhead) compared to aQDs only application and vehicle controls. Data is shown as mean ± SEM, n = 4-5 mice/group, control and aQD neutrophil counts same data as in Fig. 2B, Two-way ANOVA test, green stars indicate significances between anti-E-selectin mAb + aQDs and aQDs group. (B) Analysis of E-selectin expression. Representative lung slices from control and aQDs-treated mice after 1h, stained with rat anti-E-selectin antibody (red), Alexa488-labeled anti-Ly6G antibody (green), rabbit Anti-PRX antibody (white) and DAPI (blue). (Scale bar: 10/20 µm). (C) Quantification of E-selectin positive microvessels from immunostainings. Data is shown as mean ± SEM. n = 3 mice (6 FOV) /group, Student’s t-test. * indicates P ≤ 0.05, ** indicates P ≤ 0.01, *** indicates P ≤ 0.001, and **** indicates P ≤ 0.0001.

**Figure 5:**
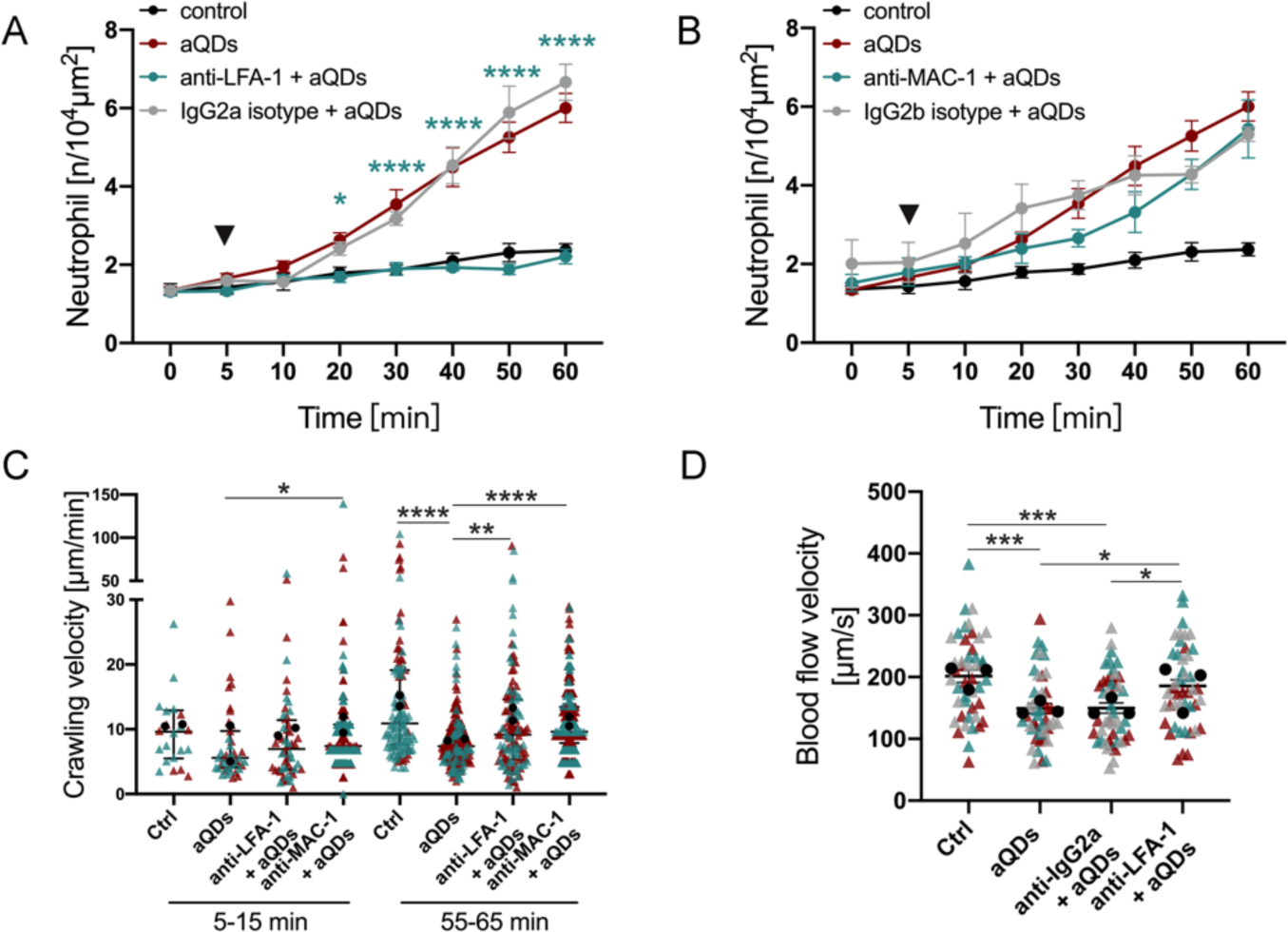
LFA-1 and MAC-1 contribute to aQD induced neutrophil recruitment. Quantification of recruited neutrophils over time in mice pre-treated intravenously with (A) anti-LFA-1 mAbs or (B) anti-MAC-1 mAbs or isotype control Abs (30 μg/mouse) 30 min prior to L-IVM followed by aQDs application at 5 min (arrowhead). Data is shown as mean ± SEM, n = 3-4 mice/group, control and aQD neutrophil counts same data as in Fig. 2B, Two-way ANOVA test, green stars indicate significances between anti-LFA-1/anti-MAC-1 mAbs + aQDs and aQDs group. (C) Velocity of crawling neutrophils under different conditions at 5-15 min and 55-65 min. Data is shown as median (IQR) and each measurement is represented by a colored triangle, n = 22-365 crawling neutrophils over 10 min from 2 mice/group, Kruskal-Wallis test. Mean values of the individual mice are shown as black dots. (D) Quantification of blood flow velocities in the different experimental groups at 1 h. Data is shown as mean ± SEM, each measurement is represented by a colored triangle, n = 45 measurements from 3 mice/group, One-way ANOVA test. Mean values of the individual mice are shown as black dots. * indicates P ≤ 0.05, ** indicates P ≤ 0.01, *** indicates P ≤ 0.001, and **** indicates P ≤ 0.0001.

### Endothelial selectins are required for neutrophil blood-vessel interactions

Neutrophil recruitment requires adhesion at blood-vessel walls, which is mediated by adhesion molecules, including selectins in most extra pulmonal tissues. At the lung endothelium, however, neutrophil binding to selectins has been reported to be stimulus dependent. (*38, 49*)

To investigate the role of selectins in aQDs-induced neutrophil recruitment within the pulmonary microcirculation, specific neutralizing antibodies targeting selectins were administered intravenously 30 min before L-IVM and aQDs injection. Treatment with anti-E-selectin antibodies prior to aQDs application resulted in, over time and relative to controls, unchanged neutrophil counts (after 60 min: 1.95 ± 0.40/10^4^μm^2^), indicating the requirement of E-selectin for aQDs-induced neutrophil recruitment (Fig. 4 A). Conversely, pretreatment with anti-P-selectin antibodies did not decrease neutrophil numbers post aQDs application (Sup. 7). Histological examination (Fig. 4 B and C) revealed a significant increase in E-selectin-positive microvessel segments after aQDs exposure compared to control lung tissue (21 ± 1.62/FOV vs. 3.33 ± 0.84/FOV), thereby emphasizing endothelial activation and E-selectin involvement in aQDs-induced neutrophil recruitment.

### LFA-1 and MAC-1 are required for neutrophil recruitment

For the inflammatory responses in the vascular bed of extrapulmonary tissues, previous IVM studies have established the essential roles of the leukocyte integrins LFA-1 (CD11a/CD18) and Mac-1 (CD11b/CD18) for adhesion, crawling and transmigration (*50–52*). However, the contribution of these integrins for neutrophil responses in NPs-induced sterile pulmonary vascular inflammation is not clear. Accordingly, mice were pretreated with anti-LFA-1 or anti-MAC-1 neutralizing antibodies prior to aQDs administration. Anti-LFA-1 antibodies effectively suppressed aQDs-induced neutrophil recruitment, maintaining neutrophil counts comparable to the control baseline (1.31 ± 0.10/10^4^µm^2^ at 0 min, increasing to 2.21 ± 0.19/10^4^µm^2^ at 60 min) (Fig. 5 A). Anti-MAC-1 antibodies, in contrast, failed to reduce neutrophil levels observed 60 min after aQDs but only delayed the response (Fig. 5 B). Blocking of MAC-1 immediately increased neutrophil crawling velocity compared to aQDs group (7.40 (3.94) µm/min vs. 5.59 (5.65) µm/min). Both anti-LFA-1 and anti-MAC-1 mAbs reversed the reduced crawling velocities induced by aQDs at 55-65 minutes (9.2 µm/min and 9.6 µm/min, respectively, vs. 7.4 µm/min), almost reaching control levels (10.9 µm/min) (Fig. 5 C). Additionally, blood flow velocities significantly increased from 149.2 ± 8.23 µm/s to 185.7 ± 9.79 µm/s after anti-LFA-1 antibody treatment, indicating that inhibiting LFA-1 not only suppressed neutrophil recruitment but also restored blood flow velocity in response to aQDs (Fig. 5 D).

### The eATP/P2X7R axis drives aQD induced pulmonary inflammation via E-selection expression

We hypothesized that aQDs in the bloodstream might induce cell stress, triggering the release of DAMPs, particularly extracellular ATP (eATP), a known pro-inflammatory stimulant associated with tissue injury and immune cell activation (*53–55*). In examining the potential involvement of eATP on immune responses following QDs treatment, plasma eATP levels were analyzed. Fig. 6 A illustrates a noteworthy rise in systemic eATP concentrations after 1 hour of aQDs application, increasing to significant levels after 24 hours. This may indicate potential cellular stress and systemic pro-inflammatory activation. P2X7R, a member of the purinergic P2 receptor family, expressed on inflammatory cells like neutrophils (*56*), monocytes (*57*), macrophages (*58, 59*), and endothelium (*60*), and known to sense eATP (*61*), serves as an ATP-gated cation channel, leading to ion flux (Ca^2+^ and Na^+^ influx, K^+^ efflux) and various subsequent cellular responses, including inflammasome activation, IL-1b release and neutrophil recruitment (*62–64*).

**Figure 6:**
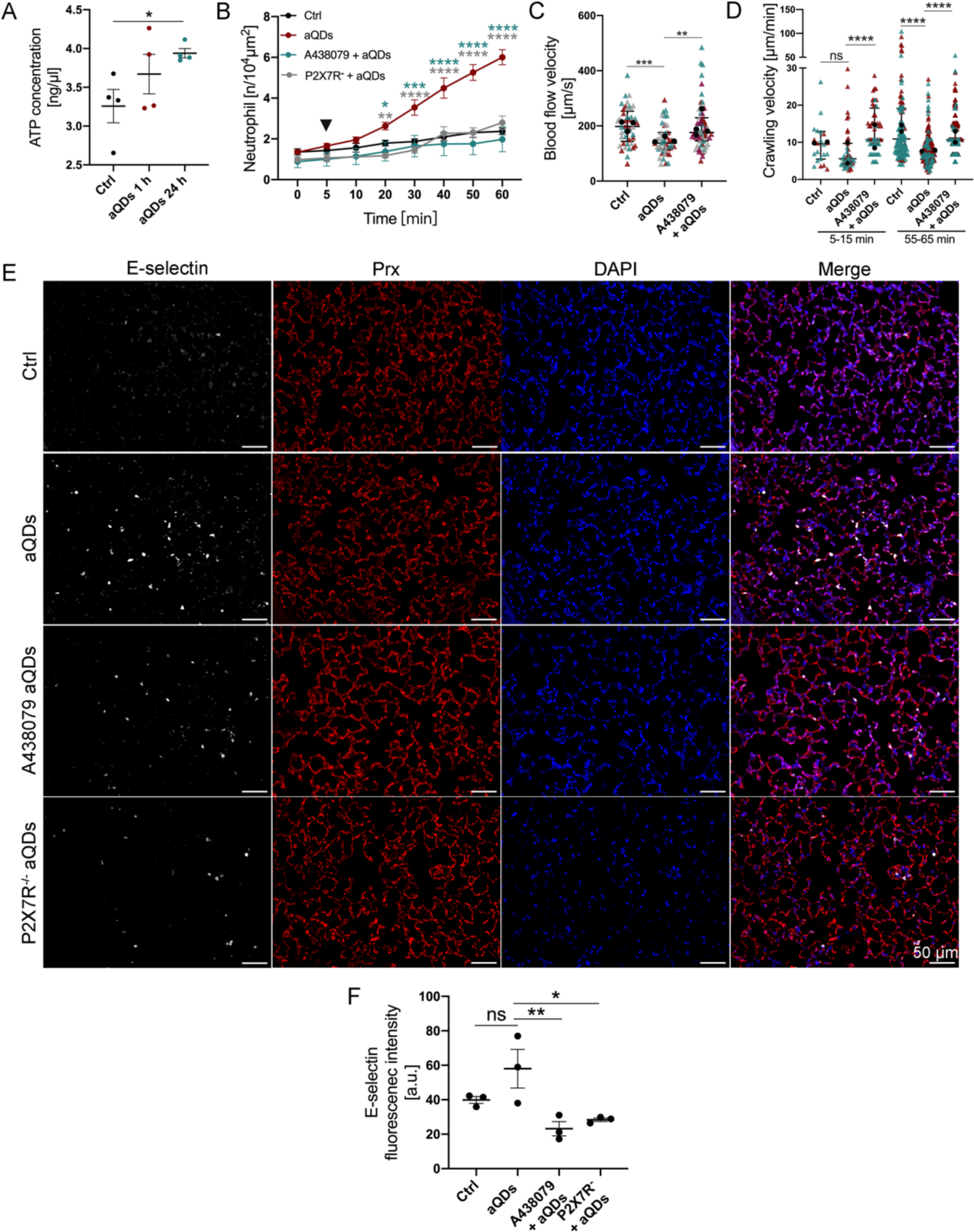
eATP/P2X7R axis is involved in aQDs-induced neutrophil recruitment. (A) Quantification of eATP concentrations in plasma samples after different treatments. Data is shown as mean ± SEM, n = 4 mice/group, Student’s t-test. (B) Quantification of recruited neutrophils over time after aQDs application (arrowhead) in mice with and without pre-treatment by intravenous injection of P2X7R antagonist (A438079, 30 μg/mouse) 30 min prior to L-IVM and in P2X7R knockout. Data is shown as mean ± SEM, n = 4-6 mice/group, control and aQD neutrophil counts same data as in Fig. 2B, Two-way ANOVA test, green stars indicate significances between P2X7R antagonist + aQDs and aQDs groups, grey stars indicate significances between P2X7R knockout mice + aQDs and WT mice + aQDs groups. (C) Quantification of blood flow velocity after 1h of L-IVM. Data is shown as median (IQR) and each measurement is represented by a colored triangle, n = 45-60 measurements from 3 mice/group, Kruskal-Wallis test. Mean values of the individual mice are shown as black dots. (D) Crawling velocities of neutrophils upon P2X7R antagonist pretreatment and aQDs application. Data is shown as median (IQR) and and each measurement is represented by a colored triangle, n = 22-247 neutrophils from 2 mice/group, Kruskal-Wallis test. Mean values of the individual mice are shown as black dots. (E) Histology of E-selectin in lung slices from control mice, aQDs-treated mice, A438079 pretreatment mice receiving aQDs, and P2X7R knockout mice with aQDs application for 1h. Rat anti-E-selectin antibody (red), rabbit Anti-PRX antibody (white) and DAPI (blue) were used. (Scale bar: 50 µm). (F) Expression of E-selectin quantified by relative fluorescence intensity from these immunostainings. Relative fluorescent intensities for E-selectin were corrected against the background of lung tissue. Data is shown as mean ± SEM (round dots), each measurement is represented by a colored triangle, n = 3 mice (6 FOV) /group, One-way ANOVA test. * indicates P ≤ 0.05, ** indicates P ≤ 0.01, *** indicates P ≤ 0.001, and **** indicates P ≤ 0.0001.

We employed the highly selective and potent P2X7R antagonist (A438079) (*65*) to investigate the impact of ATP sensing on neutrophil recruitment. Pretreatment with A438079 before aQDs injection resulted in reduced neutrophil recruitment, aligning with control levels (1.98 ± 0.61/104μm^2^ at 60 min) (Fig. 6 B). Moreover, blood flow velocity significantly increased to 175.9 μm/s (Fig. 6 C). A438079 notably ameliorated the diminished neutrophil crawling velocity induced by aQDs, raising it from 5.6 μm/min to 10.7 μm/min initially and from 7.4 μm/min to 10.7 μm/min at 60 min (Fig. 6 D). The essential contribution of P2X7R was additionally confirmed in P2X7R knock-out mice (Fig. 6 B).

Since P2X7 stimulation has been described as promoter of E-selectin expression in atherosclerosis (*66*), we investigated further whether P2X7 regulates the expression of E-selectin in the pulmonary microcirculation upon aQD triggered inflammation. E-selectin fluorescence intensities were significantly reduced in A438079 pretreated as well as in P2X7R^−/−^ lung tissue slices 1h after aQD application when compared to aQD-treated controls (Fig. 6 E and F).

These findings suggest that P2X7R inhibition holds promise in mitigating NP-induced vascular inflammation and partially restoring normal neutrophil function. This presents a potential therapeutic avenue for ameliorating the adverse effects of NPs on the pulmonary microcirculation and inflammation.

### aQD application alters systemic cytokine patterns

Building upon our findings of aQDs’ influence on the pulmonary microcirculation, we further investigate the systemic effects of QDs application. Following i.v. administration of aQDs, cytokine concentration changes in the systemic circulation were studied. Clusters of cytokine levels in the systemic circulation notably increased after 1 hour of aQDs application, partly akin to LPS stimulation (Fig. 7 A). Specifically, IL-6, CCL4, CCL17, CCL22, CCL24, and CCL27 displayed a significant increment after 1h of aQDs and 4h of LPS treatment compared to control groups, while IL-10 and CCL3 were significantly elevated in the aQDs 1h group (Fig. 7 B). The heatmap representation highlights the transient response, with cytokine levels at 24h declined to controls.

**Figure 7:**
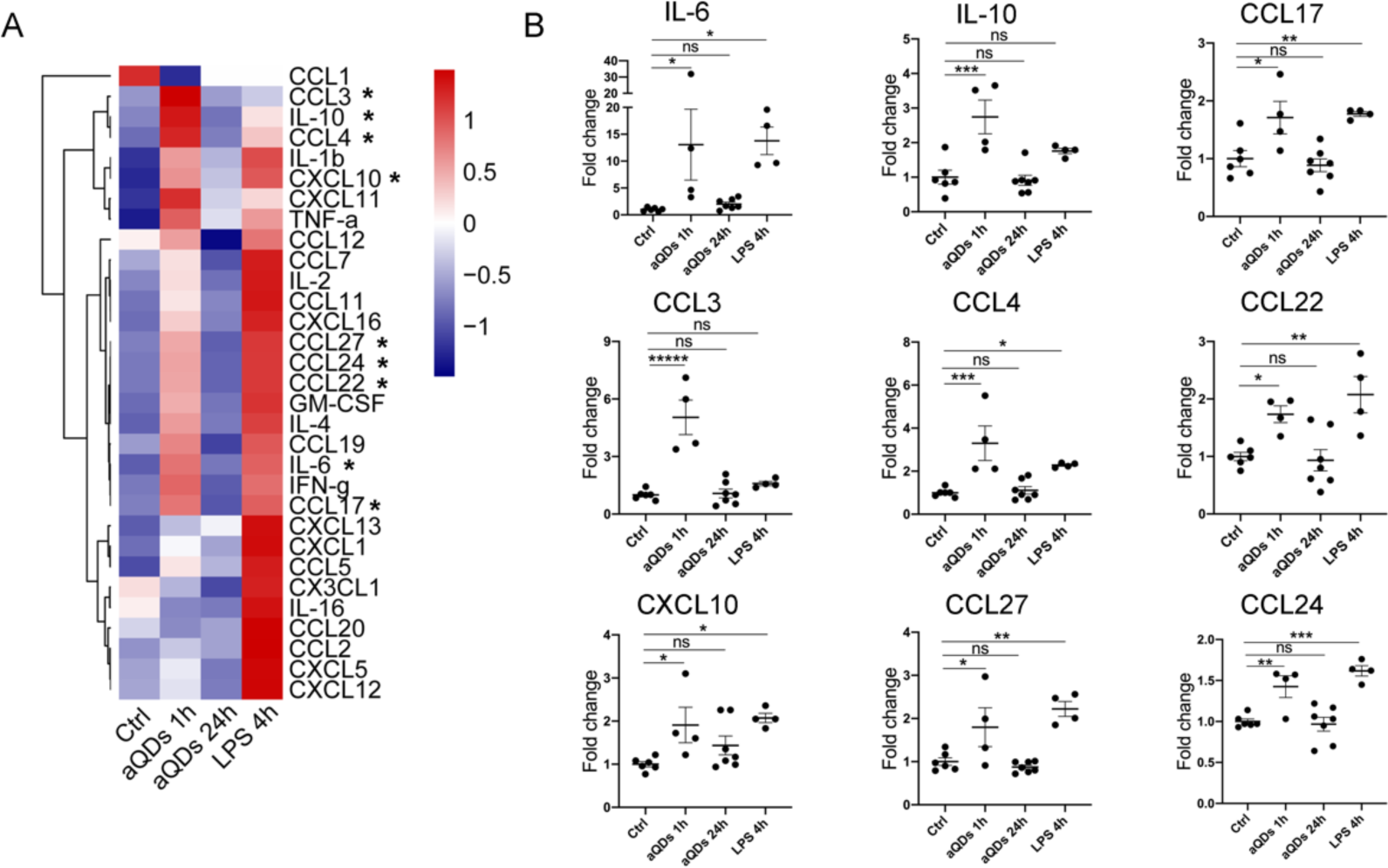
QD application induces cytokine secretion. (A) Heatmap of cytokine changes. (B) Individual alterations of cytokines. Cytokine concentrations were detected in mouse serum samples upon i.v. application of aQDs for 1h and 24h, and LPS for 4h as well as under control conditions. To combine two separate experimental rounds, the values were processed based on the mean of controls scaled to one. Data is shown as mean ± SEM, n = 4-7 mice/group, Student’s t-test, * indicates P ≤ 0.05, ** indicates P ≤ 0.01, *** indicates P ≤ 0.001, and **** indicates P ≤ 0.0001.

Taken together, our study demonstrates, that aQDs-NPs induced rapidly sterile inflammation / neutrophil recruitment in the pulmonary microcirculation. Mechanistically, this involves cellular degranulation and TNF-α release and depends on the eATP/P2X7R axis which in turn affects the expression of E-selectin. The cascade of neutrophil recruitment is further associated with LFA-1 and MAC-1. For nanomedicines, targeting the involved integrin or P2X7 receptors might present a therapeutic avenue to mitigate transient, inflammatory side effects in the pulmonary microcirculation.

## Discussion

Engineered NPs hold great promise for applications in drug delivery, diagnostics, and biomedical imaging (*67, 68*). Diverse modifications of NPs facilitate active, passive, or physicochemical targeting to specific sites, including the pulmonary endothelium, neutrophils, or areas of inflammation in conditions like acute respiratory distress syndrome, pulmonary arterial hypertension, and lung cancer, enabling effective delivery of therapeutic agents to the lungs (*69–72*). Despite the potential benefits of nanomedicines targeting the lungs, there are still risks of unintended side effects as it is the case for involuntarily inhaled nanoparticles (*2, 73–75*). Neutrophils in the pulmonary microcirculation, considered as the marginated pool, actively patrol and defend against pathogens to sustain immune defense, a role typically dominated by macrophages in extra-pulmonal tissues (*12–14, 42*). Current literature primarily focuses on the most prominent phase of the immune responses, i.e. at the response peak which occurs at least several hours after application triggered by the respective NPs (*76–79*). Here, we delved into the intricate molecular events of sterile particle-triggered vascular inflammation in the lungs, focusing on the initial steps and mechanisms of neutrophil involvement. Employing L-IVM, we captured real-time neutrophil interactions post NP challenge in the pulmonary vasculature. We show that intravenous aQD administration induced early eATP release, P2X7 receptor activation and neutrophil recruitment. This cascade involved E-selctin expression on endothelium and neutrophil LFA-1/MAC-1 function.

The QDs used in our study are either directly functionalized with carboxyl residues or coated with amino derivatized polyethylene glycol (PEG). Pharmaceutical coatings with PEG, a biocompatible polymer, enhances circulation time, by reducing interactions of plasma proteins and immune cells, and preventing renal excretion. Therefore, it is widely applied in drug delivery and biomedical research (*80, 81*). Own previous research in the extrapulmonary microcirculation showed that under physiological conditions, cQDs, but not aQDs, induced leukocyte recruitment. This was mediated by uptake of cQDs into perivascular macrophages from the blood stream, which lead to macrophage and mast cell activation, thereby triggering the canonical leukocyte recruitment cascade (*33*). In the present study, we show that in the pulmonary microcirculation aQDs triggered immediate neutrophil recruitment already 20 min after application, while cQDs induced only a slight neutrophil response. Together with increased circulating neutrophil numbers, presumably released from the bone marrow and likely mediated by elevated cytokines like CCL3/CCL4 and IL-6 in the systemic circulation, we provide evidence that aQDs trigger recruitment of additional neutrophils from both the bone marrow and the circulating blood. In addition, the influx of neutrophils into the alveolar space, which was also observed after 24 hours, and elevated eATP concentrations in the circulation also suggest a prolonged effect of aQDs in the lung. The active involvement of neutrophils in response to aQDs suggests a specific pulmonary immune response, potentially due to NP-induced stress or injury.

NPs may cause cell death through mechanisms like oxidative stress, inflammation, and membrane damage, while even low doses can disrupt cell cycle and impair cellular functions (*82–84*). Particularly under sterile, pathogen free conditions, the immune system can be activated by damage-associated molecular patterns (DAMPs), including eATP released from injured or damaged cells and tissue. Extracellular ATP initiates an immune cascade involving sentinel cells like mast cells, macrophages, and dendritic cells upon release from injured or damaged tissue (*54, 53, 55*) significantly promotes neutrophil migration during LPS-induced inflammation (*85*). P2X7R, an ATP-gated cation channel is highly expressed on multiple immune cells (*56–59, 86*) and endothelial cells (*60*). However, controversy exists regarding P2X7R expression in neutrophils, which has been accepted although certain studies have shown the contrary (*86, 87, 56*). The P2X7R channel is activated by high concentrations of eATP and allows Na^+^ and Ca^2+^ influx and K^+^ efflux. Repeated or prolonged stimulation induces formation of a non-selective pore, culminating in cytotoxic activity and cell death (*88–90*). P2X7R has been shown to be involved in cellular migration (microglia) and proliferation (*91*). Ion flux correlates with neutrophil activation and recruitment (*62, 92–95*). Our findings demonstrated that P2X7R inhibition/knockout abolished neutrophil recruitment elicited by aQDs. We cannot exclude that inhibition of P2X7R directly affects neutrophils (*96*), modifying their normal cellular responses to eATP and thereby modulating neutrophil crawling and transmigration. However, more likely P2X7R on endothelial cells is implicated in triggering E-selectin expression, in agreement with studies (*66*), and consequently recruitment of neutrophils in the lungs.

In addition, ATP released by vascular immune cells, and/or endothelial cells might activate P2X7R, thus contributing to the release of proinflammatory mediators by macrophages. These mediators, in turn, activate endothelial cells and neutrophils in a positive feedback loop (*97*). Posttranslational upregulation of E-selectin facilitates very rapid E-selectin cell surface expression (*98*). In summary, our finding that P2X7R inhibition blocks neutrophil recruitment demonstrates a key role for P2X7R in NP-induced inflammation in the lung and thus offers a potential point of application for therapeutic interventions and to mitigate side effects triggered by therapeutic NPs.

The initiation and regulation of acute pulmonary inflammation involve various cytokines and chemokines. Among these, key cytokines like TNF-α and IL-6 play crucial roles in early systemic inflammation and acute phase response to facilitate neutrophil recruitment, while IL-6 regulates neutrophil trafficking and prompts their migration from the bone marrow to the lungs (*99, 100*). MIP-1α/CCL3, and MIP-1β/CCL4 are chemotactic and proinflammatory regulators expressed by lymphocytes, monocytes, macrophages, neutrophils, and injured endothelial/epithelial cells (*101*). During inflammation, both CCL3 and CCL4 can contribute to the expression of proinflammatory cytokines like IL-6 to mediate neutrophil recruitment in response to inflammatory stimuli (*102, 103*). Leukocyte sequestration in the lungs, impacting microvascular flow (*104*), was evident with aQDs, causing inflammation and reduced blood flow, partially reversed by TNF-α blockage, LFA-1 inhibition, and P2X7R antagonism. Due to the extensively parallel nature of the capillary bed, neutrophil accumulation seldom leads to full occlusion of pulmonary vascular blood flow. Neutrophils engage in tethering, adhesion, and crawling processes, allowing interactions with endothelial cells and surveillance for potential pathogens (*42, 105*). This neutrophil behavior was also observed in the present study. Recruited neutrophils tend to accumulate in small capillaries that interconnect the outsides of alveoli (*15, 42, 106, 107*). Our results showed that neutrophils recruited in response to aQDs are localized predominantly in vessels smaller than 20 μm. The onset of neutrophil recruitment is mediated by multiple selectins and integrins. Contradictory findings exist regarding the role of E-, P-, and L-selectins in neutrophil recruitment in the lung, which seems to depend on the specific inflammatory stimulus in the lung (*15, 55, 107–110*). We observed that inhibiting E-selectin but not P-selectin effectively reduced the recruitment of neutrophils. Since leukocyte rolling does not occur in the pulmonary microcirculation (*12, 55*), P-selectin might be dispensable.

Our findings highlight the distinctive response of neutrophils in the pulmonary microcirculation to different types of QDs, emphasizing the potency of NPs to induce rapid neutrophil recruitment in the lung, thus extending the current understanding of sterile particle-triggered inflammation. Crucially involved are the release of TNF-α and eATP, activation of the P2X7R, and subsequent expression of E-selectin and involvement LFA-1, and MAC-1. By inhibiting each of these key players, we could modulate neutrophil dynamics, and thereby mitigate the inflammatory, adverse effects of NP exposure. Together with the observed reduction of pulmonary microvascular blood flow reversible by the various interventions, this highlights potential avenues for therapeutic interventions or even reductions of side effects from therapeutic/theranostic NPs.

The divergent impacts of QDs with distinct surface modifications and compositions in different vascular beds, as elucidated in this study and previous own studies (*20, 24, 33, 111*), underscore the importance of their thorough evaluation in the design and development of nanomedicines. The careful consideration of NP characteristics, such as their impact on immune responses and vascular inflammation, is critical for ensuring the safety and efficacy of nanomedical applications.

In conclusion, our comprehensive analysis unveils novel insights into the initial stages of sterile particle-elicited vascular inflammation in the lung. Specifically, we elucidate that eATP-gated P2X7R serves as a trigger, initiating endothelial activation and subsequent neutrophil recruitment. This knowledge not only advances our understanding of pulmonary immune responses to NPs but also offers a foundation for developing safer nanomedicines and addressing the side effects associated with pulmonary vascular inflammation.

## Methods

### Animals

Female C57BL/6N mice (8-12 weeks old) were purchased from Charles River (Sulzfeld, Germany). Breeding of C57BL/6N-*P2rx7*^t^ ^m1d(EUCOMM)Wtsi^ (P2X7^−/−^) mice from *P2rx7* ^tm1a(EUCOMM)Wtsi^ mice has been described (*112*). Mice were housed in individually ventilated cages with free access to food and water. All animal experiments followed strict ethical guidelines and were approved by the local Animal Care and Use Committee (District Government of Upper Bavaria), complying with EU Directive 2010/63/EU.

### Nanoparticles

Quantum Dots Qdot^TM^ 655 ITK^TM^ Carboxyl and Qdot^TM^ 655 ITK^TM^ Amine (PEG) Quantum Dots (QDs) were procured from Invitrogen Corporation (Karlsruhe, Germany) with an emission wavelength of 655 nm. These QDs feature a CdSe core, a ZnS shell, and either amine residues with a polyethylene glycol coating (aQDs) or carboxyl residues without coating (cQDs). The hydrodynamic diameter of aQDs in DPBS is 25.2 nm, while cQDs measure 18.1 nm (*23*). QDs were resuspended in 50 μl of DPBS at a concentration of 1 pmol/g and intravenously injected into mice through the angular vein using a 1 ml insulin syringe (Becton, Dickinson and Company, Franklin Lakes, USA). Melamine resin particles (MF-FluoRed, MFs) (ex: 636 nm, em: 686 nm, 0.94 ± 0.05 μm) were procured from microParticles GmbH (Berlin, Germany). A stock solution of MFs at 2.5% concentration was diluted with ultrapure H_2_O (GibcoTM life technologies Thermo Fisher Scientific Carlsbad, USA) to create a 0.05% working solution, which was thoroughly vortexed and intravenously injected in a volume of 50 μl for blood velocity measurements

### In vivo application of fluorescent antibodies and blocking reagents

Deep anesthesia was induced using a triple compound of medetomidine (0.5 mg/kg body weight), midazolam (5 mg/kg body weight), and fentanyl (0.05 mg/kg body weight) (MMF) by intraperitoneal injection. Fluorescent antibodies, blocking antibodies, and antagonists (Table S1) were administered intravenously 30 min prior to L-IVM, appropriately proportioned and diluted in DPBS to achieve a total volume of 80 μl.

### Lung intravital microscopy

The surgical protocol closely adhered to the details outlined in a prior publication (*30*). Deep anesthesia was induced using a triple compound of medetomidine (0.5 mg/kg body weight), midazolam (5 mg/kg body weight), and fentanyl (0.05 mg/kg body weight) (MMF) by intraperitoneal injection. Surgical sites (neck and left chest) were locally anesthetized with Bucain (50µg/site, Puren Pharma, Germany). Mice were positioned on a temperature-controlled heating pad to maintain the core body temperature at 37°C. After the initial MMF administration, a half-dose was injected every 45 min during L-IVM. Fluorescent antibodies and relevant inhibitors were administrated by i.v. injection 30 min before the L-IVM procedure. A tracheostomy was performed to insert a small catheter linked to a miniature rodent ventilator (MiniVent, Harvard Apparatus, Massachusetts, USA). The ventilator maintained 10 μl/g BW for stroke volume and 150 breaths/min for breathing rate, coupled with a positive end-expiratory pressure of 2∼3 cm H_2_O, all administered with 100% oxygen throughout the experiments.

The mice were then placed in right lateral decubitus position and a custom made flanged thoracic suction window, furnished with an 8 mm glass coverslip (VWR, Radnor, USA) was inserted into a 5 mm intercostal incision through the parietal pleura between ribs 3 and 4 of the left chest. 20–25 mmHg of suction was used to immobilize the lung by a custom-made system consisting of a differential pressure gauge (Magnehelic, Dwyer Instruments, nc, USA) and a negative pressure pump (Nupro, St Willoughby, USA). Observations were facilitated using a VisiScope. A1 imaging system (Visitron Systems GmbH, Puchheim, Germany), equipped with a water dipping objective (20x, NA 1.0, Zeiss Micro Imaging GmbH, Jena, Germany), and a 16-color LED light source for fluorescence epi-illumination (pE-4000; CoolLED, UK). Throughout the experiment, a quadband filter set was used (F66-014, DAPI/FITC/Cy3/Cy5 Quad LED ET Set; AHF Analysentechnik AG, Tuebingen, Germany). The 470 nm LED module was employed at 40% output power for 50 ms to excite QDs and Alexa488-labeled anti-Ly6G antibody. Emissions from QDs and antibodies were separated by a beam splitter (T 580 lpxxr, Chroma Technology Corp, Bellows Falls, USA), then captured by two Rolera EM2 cameras and processed using VisiView software (Visitron Systems GmbH, Puchheim, Germany). After 5 minutes of observation to obtain baseline conditions, QDs at 1 pmol/g, diluted with DPBS to a total of 50 μl, were i.v. administered.

### Quantification of neutrophil numbers and blood flow velocity

After the thoracic surgery, the L-IVM imaging in mice was recorded, marking time point 0. Five random fields of view were captured within the imaging window every 5-10 min for QDs and neutrophils in each mouse. Each imaging session typically takes 30 s. The images were calibrated to a scale of 2.5 pixels/μm using Fiji software (Rasband, W.S., U. S. National Institutes of Health). Neutrophil counts were assessed using the “Trackmate” plugin (*113*) in Fiji, with an estimated diameter of 10 μm. For precise measurement of neutrophil migration parameters in mouse lungs, obtaining high-quality images without excessive shaking was crucial. Five random regions were identified at 0 minutes to assess imaging quality. From 5 min onward, images of one region were captured every 5 seconds. The image displacement was rectified using the “Register virtual stack slices” plugin (*114*) in Fiji, employing the “Translation” feature extraction and registration models. Subsequently, the velocity and displacement of neutrophils were analyzed using the “Trackmate” function in Fiji. Following 1 hour of L-IVM, MFs particles for blood flow measurements were intravenously introduced, and images were captured at intervals of 0.185 s. These sequential images underwent calibration to a scale of 2.5 pixels/μm. Blood flow velocity was ascertained by monitoring the trajectory of MF fluorescence particles using “Manual Tracking” plugin.

### Bronchoalveolar lavage (BAL) preparation and cell differentiation

Upon completion of L-IVM imaging, deeply anesthetized mice were humanely euthanized by abdominal aorta exsanguination. DPBS, instilled into the lungs through the tracheal cannula and aspirated by pulling on the syringe barrel, was used for bronchoalveolar lavage fluid (BALF) collection. The first 2 ml and subsequent 8 ml of recovered BALF were collected separately and kept on ice. After centrifugation (4°C, 20 min, 400 g), the first 2 ml of supernatant was aliquoted and stored at −80°C for cytokine analysis. The cell pellet obtained from the entire 10 ml of BALF was collected and counted with 0.2 % Trypan blue dye (Sigma Aldrich, Taufkirchen, Germany). For each cytospin, 3 × 10^4 BAL cells were utilized. Cytospin slides were stained with May-Grünwald-Giemsa staining (Sigma Aldrich, Taufkirchen, Germany) to identify macrophages, neutrophils, lymphocytes, and monocytes.

### Light-sheet fluorescence microscopy (LSFM)

The mouse lungs underwent perfusion with DPBS and were fixed overnight in 4% PFA. Optical clearance was achieved following a previously published protocol (*115*). Shortly, the fixed tissue underwent a series of incubations in Tetrahydrofuran (THF, Sigma) at varying concentrations, followed by immersion in Dichloromethane (DCM, Sigma) and dibenzyl ether (DBE, Sigma). Imaging was conducted using a light sheet fluorescence microscope (LSFM, Ultramicroscope II, LaVision Biotec). QDs were excited at 470 ± 30 nm, and emission was detected at 640 ± 30 nm, while lung autofluorescence was excited at 520 ± 40 nm and detected at 585 ± 40 nm. LSFM images were acquired and processed using Imaris 9.1.0 software (Bitplane, Belfast, United Kingdom).

### Blood analysis

One hour post i.v. administration of QDs (1 pmol/g), mice were anesthetized with xylazine and ketamine (i.p.). Subsequently, 50 μl of blood was collected from the angular vein using a microcapillary pipette (Hirschmann Minicaps, Laborgeräte GmbH & Co. KG, Germany) and diluted 1:7 with Cellpack DCL buffer provided by Sysmex Deutschland GmbH. A complete hematology analysis, encompassing white blood cell (WBC) count, platelet count, erythrocyte-related parameters, and percentages of different WBC types, was conducted using a Hematology Analyzer (Sysmex XN-1000V) using the capillary mode with mouse settings provided by the instrument.

### Bio-Plex Pro Mouse Chemokine Panel 31-Plex Assay

Blood and BALF samples were collected from mice after 1 hour or 24 hours of i.v. QD administration (1 pmol/g) and after intratracheal instillation of LPS for 4 hours. Cytokines and chemokines in the blood and BALF samples were quantified following the manufacturer’s protocol for the Bio-Plex Pro Mouse Chemokine Panel 31-Plex (Bio-Rad Laboratories, California, USA). Data acquisition was performed using a LuminexTM 200 plate reader (InvitrogenTM, Thermo Fisher Scientific, Carlsbad, USA), and the BioPlex Manager 6.2 software (Bio-Rad Laboratories, California, USA) was used for analysis. Standard curves were fitted using the logistic-5PL regression type. A heatmap based on relative abundancies of cytokine concentrations was generated using “z score” and “pheatmap” packages (Kolde R, 2019, R package version 1.0.12) in R software (R-4.2.3).

### Toluidine blue staining

Toluidine Blue (C.I.52040, Merck Millipore, Germany) was used to prepare a working solution with a pH range of 2.0-2.5 following the manufacturer’s instructions. Lung slices were deparaffinized by sequential immersion in Xylene, 100% Ethanol, 90% Ethanol, 80% Ethanol, and 70% Ethanol, followed by rinsing with distilled water, and then were immersed in the Toluidine Blue working solution for 2 min. After a 2-min wash with distilled water, the stained slices underwent dehydration and were then mounted with coverslips. This staining technique was employed to identify mast cells, characterized by violet/red-purple metachromatic granules against a blue background tissue.

### Immunofluorescence staining

Lung slices, following deparaffinization, underwent heat-induced epitope retrieval (HIER) by incubating in citrate pH = 6.0 buffer. Subsequently, the slices were treated with a blocking solution (5 % goat serum and 0.3 % TritonX-100 (Sigma-Aldrich GmbH) in PBS) for 1 hour, followed by incubation with primary antibodies (E-selectin (ab2497; Abcam, Cambridge, UK), CD11b (ab133357; Abcam, Cambridge, UK), Prx (HPA001868; Sigma-Aldrich, Merck KGaA, Germany)) in antibody diluent (1% Bovine serum albumin and 0.3% Triton X-100 in PBS) at 4 °C overnight. Next, the slices were incubated with secondary antibodies (Goat anti-rat IgG Alexa Fluor 568, A2110844; Goat anti-rabbit IgG Alexa Fluor 568, A2155282; Thermo Fisher Scientific, Waltham, MA, USA; Goat anti-rat IgG Alexa Fluor 647, A1921562, Thermo Fisher Scientific, Waltham, MA, USA), Alexa488-labeled anti-Ly6G antibody, and DAPI (D9564; Roche, Basel, Switzerland) in antibody diluent for 1 hour. Each step was followed by a washing step in DPBS solution containing 0.1% Tween®20 for 3 times, each for 5 min. Finally, the stained slices were mounted using DAKO fluorescence mounting medium (Dako Omnis, Agilent, Santa Clara, USA) and covered with coverslips.

### Statistical analysis

All data were presented as mean ± SEM or median (IQR) and plotted using GraphPad Prism 8 (GraphPad Software Inc., La Jolla, USA). For normally distributed data, a two-sided Student’s t-test or One-way/Two-way ANOVA test was used for comparison between two or more groups, while for non-normally distributed data, the Mann-Whitney rank-sum or Kruskal-Wallis test was used. Significances levels were defined as follows: P < 0.05 (*), P < 0.01 (**), P < 0.001 (***), and P < 0.0001 (****), while P ≥ 0.05 were considered not significant (ns).

## Acknowledgements

The authors would like to particularly thank David Kutsche for his excellent technical work supporting this project. We are grateful to the members of the animal facility of Helmholtz Munich for their professional support and assistance; Dr. Lin Yang and Dr. Otmar Schmid (Institute of Lung Health and Immunity, Helmholtz Zentrum München) for support with particles and staining; Barbara Mosetter (Immunanalytik-Tissue Control of Immunocytes, IMA-TCI, Deutsches Forschungszentrum für Gesundheit und Umwelt, Helmholtz Zentrum München, Munich, Germany) for technical assistance with the Bioplex assay; Dr. Annette Feuchtinger (Research Unit Analytical Pathology, Helmholtz Munich) for support in light sheet imaging.

## Author contributions

CL and MR designed and planned the entire study. CL performed all the experiments and analyzed the data. QL provided support on animal experiments and analysis. LH provided support on analysis and figure revision. RI, BR, MHdA performed and supervised blood analysis. AN, JS and LC provided critical tools and experimental methods. CL and MR wrote the manuscript. MR, TS, and MS revised the manuscript. All authors read, discussed, improved, and approved the paper. Correspondence to MR.

## Supplemental Information

### Supplementary figures

**Supplemental figure and video 1:**
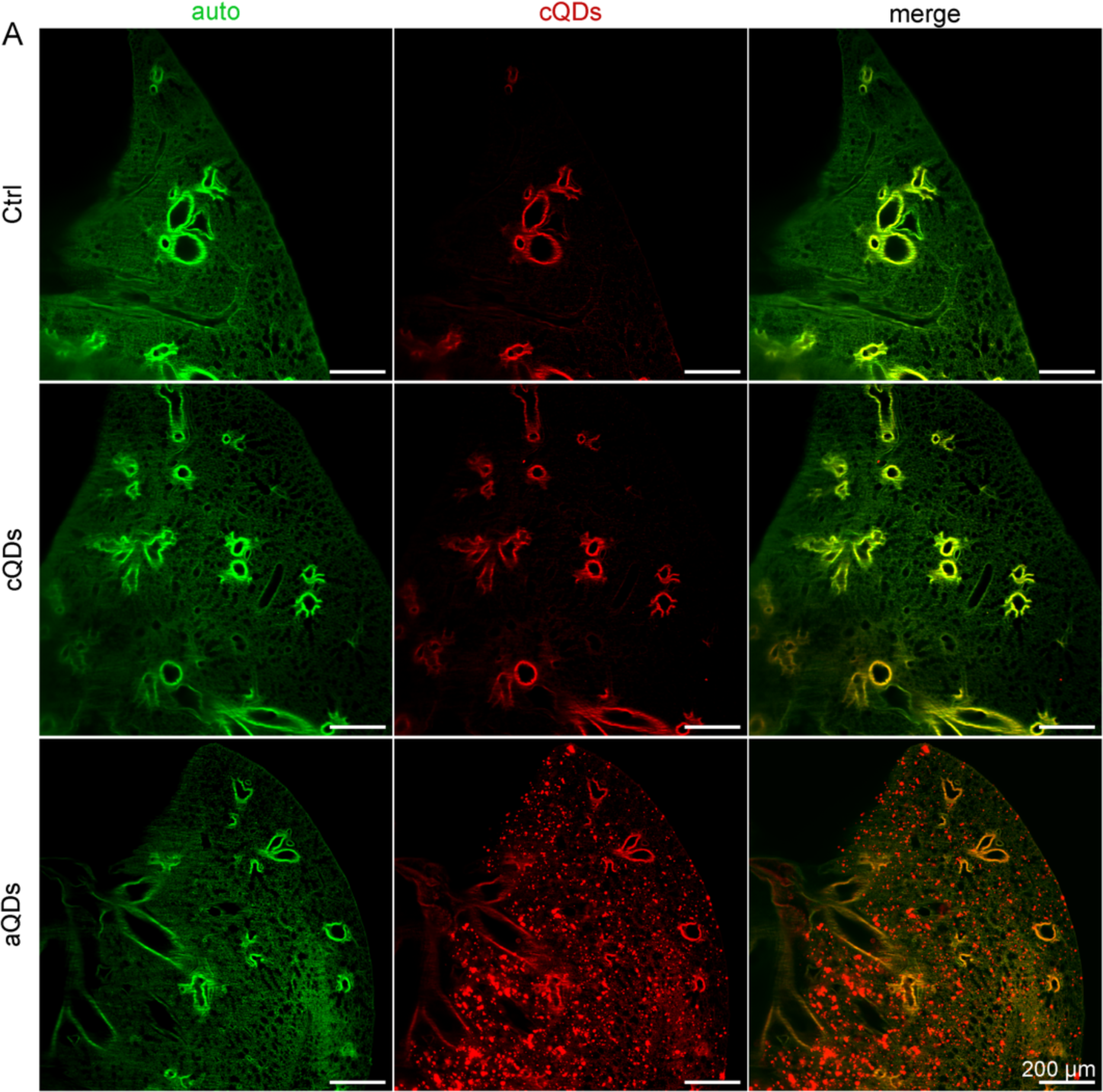
QDs distribution in whole lungs. After intravenous application of vehicle, cQDs, or aQDs for 1 hour, lung tissues were optically cleared and imaged in 3D. The lung structure is represented in green, while QDs are visualized in a bright red color, providing an overall presentation of their distribution within the lung.

**Supplemental figure 2:**
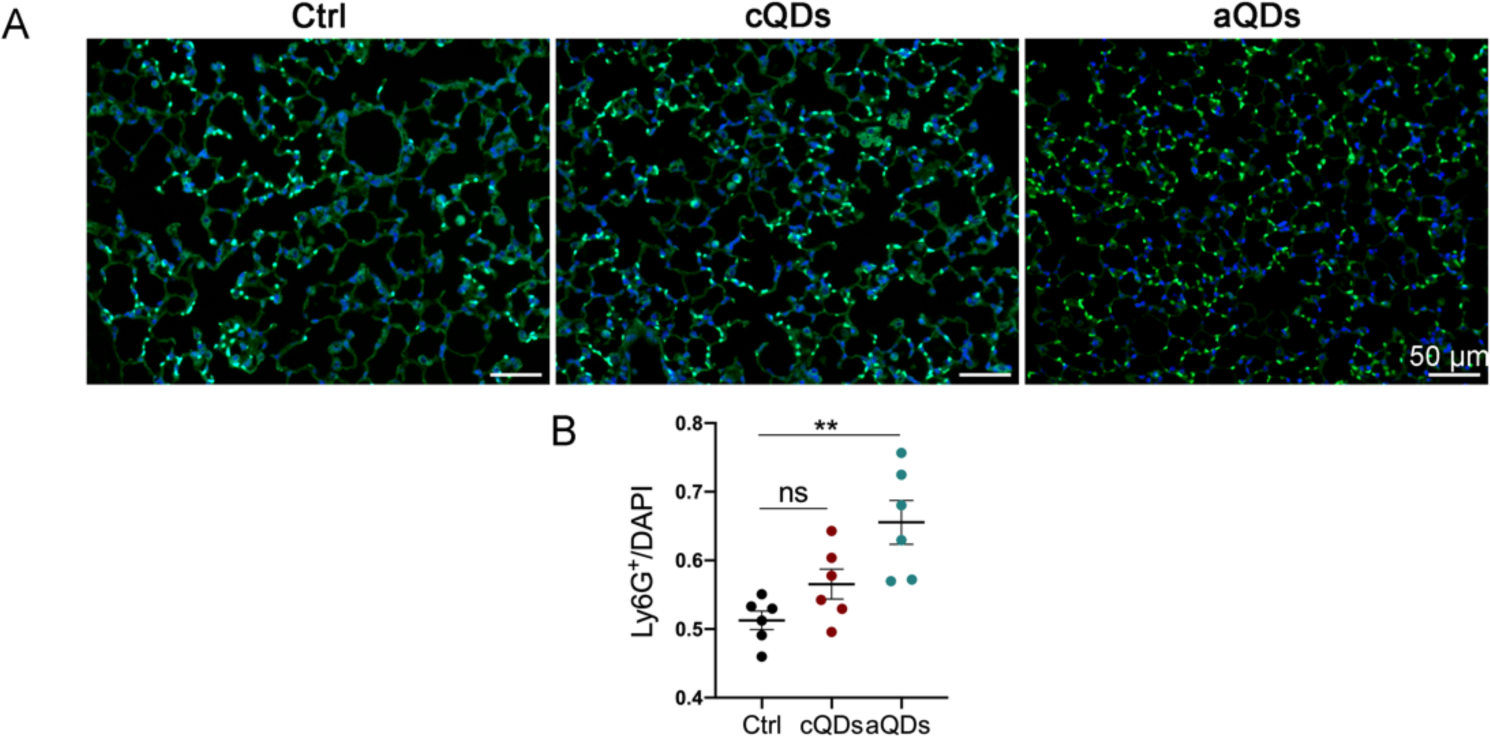
Histology of lung slices indicates increased neutrophil numbers after aQDs application. (A) Neutrophils were stained with Alexa488-labeled anti-Ly6G antibody in lung slices and are depicted in white. (B) Quantification of neutrophils is shown. n = 2 tissue slices from 3 mice/group, mean ± SEM, analyzed using Student’s t-test. * indicates P ≤ 0.05, ** indicates P ≤ 0.01, *** indicates P ≤ 0.001, and **** indicates P ≤ 0.0001.

**Supplemental figure 3:**
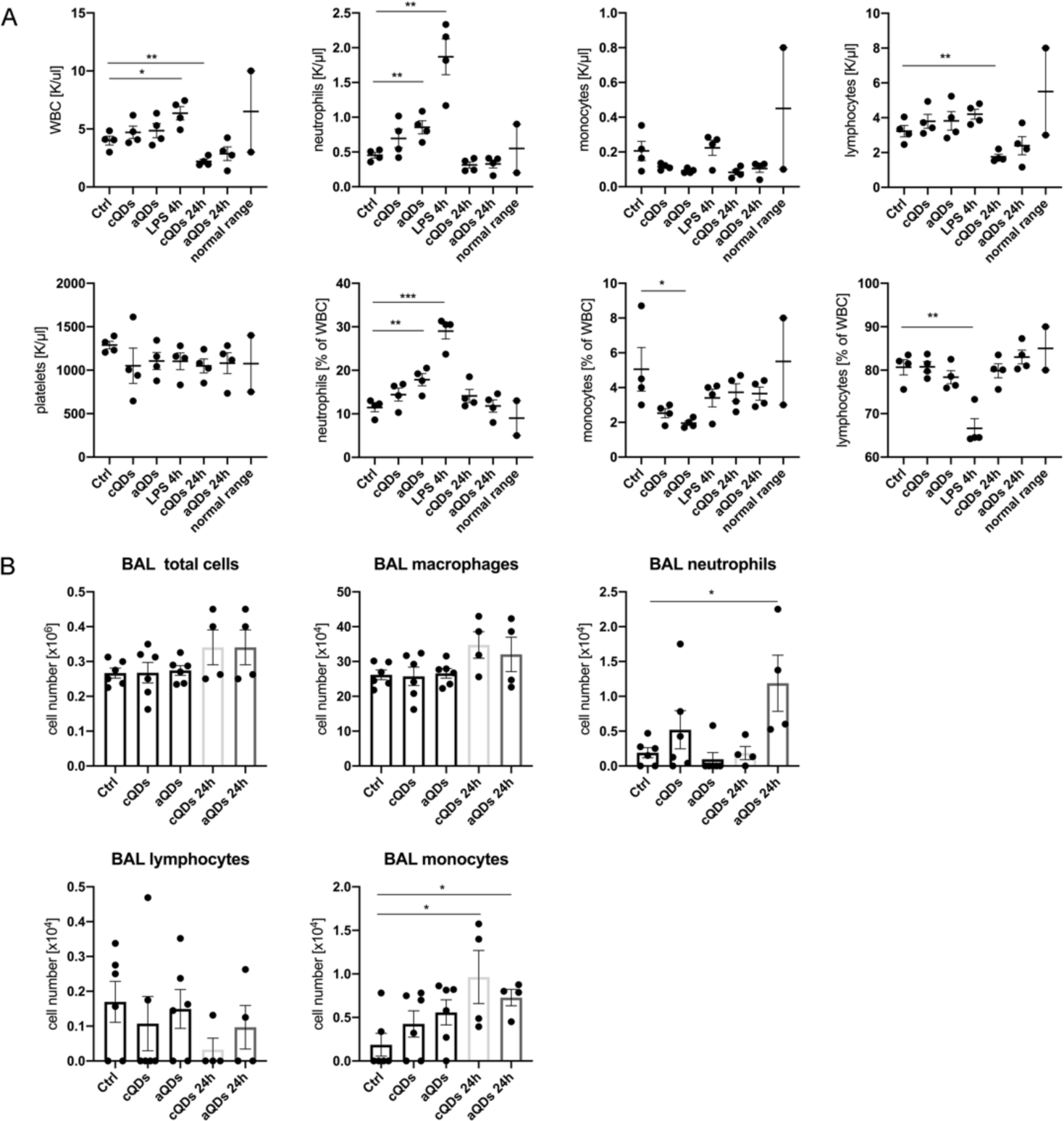
Changes in systemic blood parameters and BAL cell counts following QDs application. (A) Systemic level and percentage of immune cells 1-hour or 24-hour after QDs application. (mean ± SEM, n = 4 mice/group, Student’s t-test). (B) Quantification of total BAL cells, macrophages, neutrophils, lymphocytes, and monocytes after 1-hour or 24-hour QDs application is shown. May Grunwald-Giemsa stained cytospin samples of BAL cells were analyzed. mean ± SEM, n = 4-6 mice/group, Student’s t-test. * indicates P ≤ 0.05, ** indicates P ≤ 0.01, *** indicates P ≤ 0.001, and **** indicates P ≤ 0.0001.

**Supplemental video 4:**
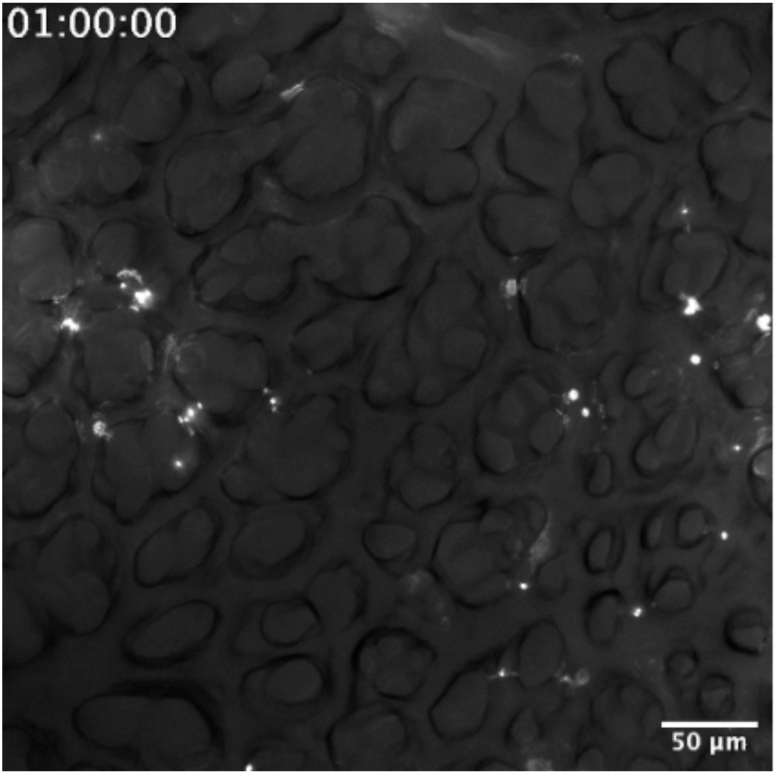
Blood flow velocity in pulmonary microvessels. Video shows blood flow depicted by bead trajectories in the lung microcirculation. The sequential images reveal blood flow direction in blood vessels presented by individual microbead trajectories, depicted as different colored lines.

**Supplemental video 5:**
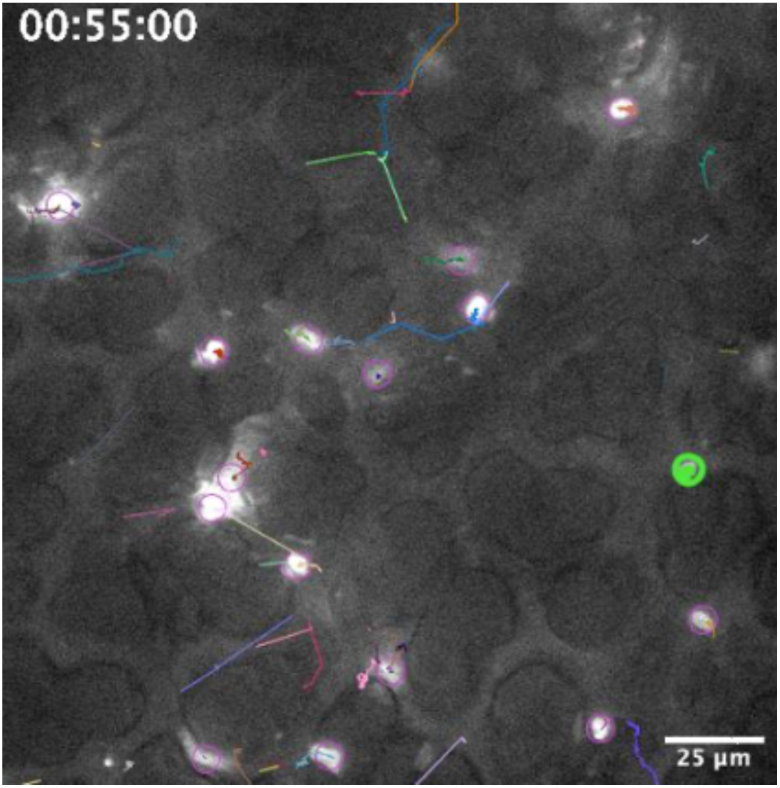
Trajectories of neutrophil movements in healthy mice via L-IVM. Neutrophil dynamics were recorded every 5 seconds over a period of 10 min by L-IVM. The images were analyzed, and the movement trajectories of neutrophils were automatically generated by plugin “Trackmate” of Fiji software. The video displays representative neutrophil trajectories during the period of 55-65 min under healthy conditions. Each trajectory is represented by a different color, indicating tracks of individual neutrophils. (Scale bar: 25 μm).

**Supplemental figure 6:**
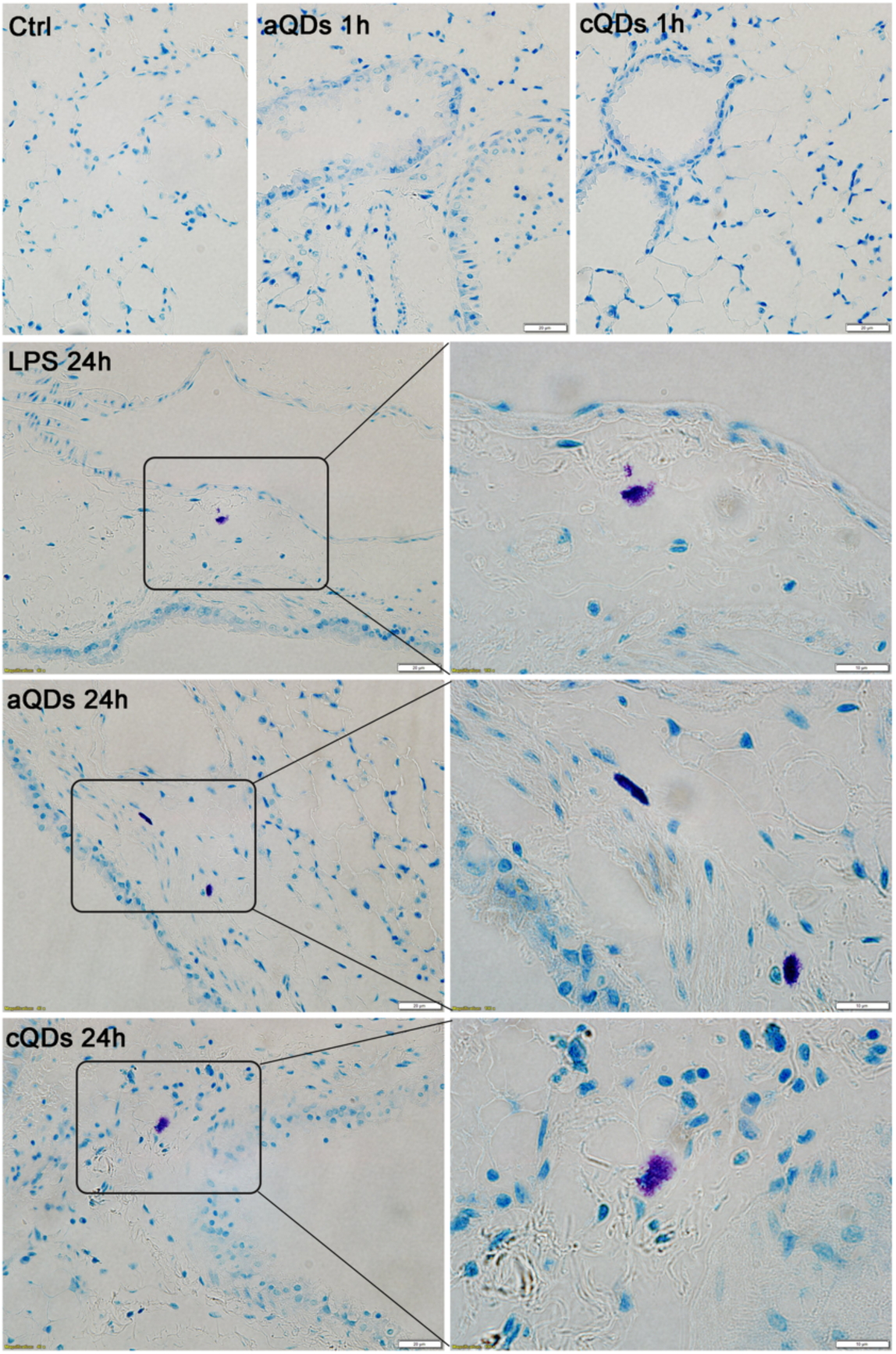
Metachromatic granule staining in mast cells after aQDs or cQDs exposure, LPS treatment, and under control conditions. The images display metachromatic granule staining in mast cells, visible as violet against the blue background of surrounding tissues, obtained after tolouidin staining, after 1 h and 24 h aQDs or cQDs exposure, 24 h LPS instillation (0.1 μg/mouse), or vehicle control. (Scale bar: 10/20 µm).

**Supplemental figure 7:**
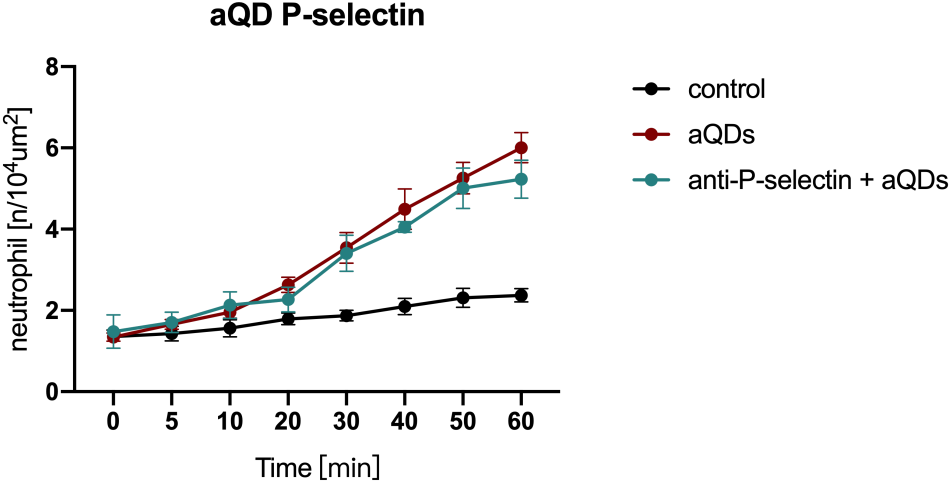
Inhibiting P-selectins did not alter neutrophil recruitment dynamics in response to aQDs. Mice were intravenously pre-treated with anti-P-selectin mAbs for 30 minutes before aQDs application, compared to aQDs-only application and control groups. Neutrophil numbers were quantified over time. Black arrow indicates aQDs injection at t = 5 min; mean ± SEM, n = 4 mice/group. Control and aQD neutrophil counts same data as in Fig. 2B.

### Supplementary table

**Supplemental Table S1:** List of in vivo fluorescent dyes, blocking antibodies, and antagonists.

| Name | Clone | Company | Dose |
| --- | --- | --- | --- |
| A438079 hydrochloride<br>(Competitive<br>P2X<sub>7</sub> antagonist) | $\geq 98$ %<br>(HPLC) | Bio-Techne GmbH,<br>Germany | 30 μg/mouse |
| Alexa Fluor® 488 anti-mouse Ly-6G antibody | 1A8 | BioLegend, Fell, Germany | 3 μg/mouse |
| Cromolyn sodium salt | assay $\geq 95$ % | Sigma Aldrich, Taufkirchen,<br>Germany | 0.2 μg/g (BW) |
| InVivoMAb anti-mouse<br>LFA-1α (CD11a) | M17/4 | Bio X Cell, Lebanon, USA | 30 μg/mouse |
| InVivoMAb anti-mouse/human CD11b | M1/70 | Bio X Cell, Lebanon, USA | 30 μg/mouse |
| InVivoMAb anti-mouse<br>E-selectin | 9A9 | Bio X Cell, Lebanon, USA | 20 μg/mouse |
| InVivoMAbs rat IgG1<br>isotype control | HRPN | Bio X Cell, Lebanon, USA | 30 μg/mouse |
| InVivoMAbs rat IgG2a<br>isotype control | 2A3 | Bio X Cell, Lebanon, USA | 30 μg/mouse |
| In vivo MAbs rat IgG2a<br>isotype control | 2A3 | Bio X Cell, Lebanon, USA | 30 μg/mouse |
| Purified anti-mouse<br>TNF-α antibody | MP6-XT22 | BioLegend, Fell, Germany | 30 μg/mouse |
| Purified rat anti-mouse<br>CD62P antibody | RB40.34 | BD Pharmingen, Germany | 20 μg/mouse |
| Ultra-LEAFTM purified<br>Rat IgG2b, κ Isotype<br>Ctrl antibody | RTK4530 | BioLegend, Fell, Germany | 20 µg/mouse |

